# Identification of conserved age-associated aggregation-prone proteins for natural molecule targeting during aging in *Caenorhabditis elegans*

**DOI:** 10.64898/2026.07.28.741274

**Authors:** Priyanka Vimal, Neha Agyal, Shagun Shagun, Shyam Kumar Masakapalli, Prasad Kasturi

## Abstract

Aging is associated with proteome remodelling and progressive accumulation of insoluble proteins. Identifying age-enriched proteins that undergo aggregation and evaluating compounds capable of modulating their behaviour may provide insights into interventions that promote healthy aging. Here, we report a proteome-guided strategy to identify age-associated aggregation-prone proteins and evaluate phytochemicals targeting conserved proteins in *Caenorhabditis elegans*. We identified proteins whose abundance increased more than four-fold in aged worms compared with young worms, many of which also accumulated in the age-associated insoluble proteome, and subsequently identified their human orthologs for comparative analysis. Based on biological relevance, structural conservation, and availability of high-confidence structural models, glutamine-fructose-6-phosphate aminotransferase-2 (GFAT-2) was selected for molecular docking. Screening of fifteen phytochemicals against *C. elegans* GFAT-2 and its human ortholog GFPT1 identified quercetin as the strongest predicted binder, exhibiting conserved interactions with both proteins. However, treatment of worms with quercetin did not significantly alter global protein insolubility during aging. This may reflect its ability to modulate inappropriate protein-protein interactions without substantially affecting the overall aggregation burden. These findings underscore the need for experimental validation of favourable in silico docking predictions. More broadly, this study provides a proteome-guided framework for prioritizing age-associated aggregation-prone proteins as candidate therapeutic targets for preserving proteostasis during aging.

## Introduction

Proteostasis, or protein homeostasis, is maintained by an intricate network of molecular chaperones, protein synthesis and degradation pathways, and quality control systems that collectively ensure correct protein folding, prevent the accumulation of damaged proteins, and preserve cellular function (Labbadia & Morimoto. 2015; Klaips et al. 2018). However, aging is accompanied by a progressive decline in proteostasis capacity due to reduced chaperone activity, impaired ubiquitin-proteasome system (UPS) function, decreased autophagic clearance, and diminished stress responses (Taylor & Dillin 2011; Hipp et al. 2019). As a consequence, damaged and misfolded proteins accumulate, promoting the formation of insoluble protein aggregates that disrupt cellular homeostasis and contribute to tissue dysfunction (Chiti F, Dobson. 2017). Protein aggregation is a hallmark of aging and underlies the pathogenesis of numerous age-associated disorders, including Alzheimer’s disease, Parkinson’s disease, Huntington’s disease, amyotrophic lateral sclerosis, and several metabolic and cardiovascular diseases (Wilson DM 3^rd^ et al. 2023) Understanding the molecular mechanisms that govern age-dependent protein aggregation and identifying strategies to modulate this process remain major goals in aging research.

*Caenorhabditis elegans* has emerged as one of the most powerful experimental systems for studying aging and proteostasis because of its short lifespan, conserved cellular pathways, availability of disease models and ease of genetic and pharmacological manipulation (Mack et al. 2018; Vimal et al. 2026). Furthermore, comprehensive age-dependent proteomic and aggregation datasets generated in *C. elegans* have enabled systematic analyses of proteome-wide changes during aging (David et al. 2010; Reis-Rodrigues et al. 2012; Walther et al. 2015; Narayan et al. 2016; Zhu et al. 2024). Several studies have demonstrated that increased protein abundance during aging is an important determinant of protein aggregation. Proteins that progressively accumulate can exceed their solubility limits, resulting in supersaturation and spontaneous aggregation even when they possess relatively low intrinsic aggregation propensity (Walther et al. 2015; Narayan et al. 2016; Zhu et al. 2024). Therefore, proteins exhibiting substantial age-dependent increases in abundance represent attractive candidates for investigating mechanisms of age-associated aggregation and for identifying therapeutic interventions capable of preventing their misfolding or aberrant interactions.

Natural phytochemicals have attracted considerable attention as potential modulators of aging and proteostasis because of their broad biological activities and favorable safety profiles (Chen et al. 2024; Cuanalo-Contreras & Moreno-Gonzalez 2019). Polyphenols and related plant-derived metabolites, including quercetin, epigallocatechin gallate, curcumin, catechins, and kaempferol, have been reported to improve stress resistance, extend lifespan in multiple model organisms, and protect against age-associated proteinopathies (Dhouafli et al. 2018; Henríquez et al. 2020; Davinelli et al. 2025). These compounds exert their beneficial effects through multiple mechanisms, including activation of stress-responsive transcription factors such as HSF-1, DAF-16/FOXO, and SKN-1/Nrf2, enhancement of antioxidant defenses, stimulation of autophagy, modulation of molecular chaperones, and inhibition of amyloid fibril formation (Wedel et al. 2018; Ren et al. 2026). Although numerous phytochemicals have shown anti-aggregation activity against disease-associated proteins such as amyloid-β, α-synuclein, and huntingtin (Freyssin et al. 2018), comparatively little is known about whether they can target endogenous proteins that naturally accumulate and aggregate during physiological aging.

Recent advances in quantitative proteomics have enabled systematic identification of proteins whose abundance changes throughout the lifespan of *C. elegans*. In particular, the comprehensive proteomic study by Walther et al. (2015) identified hundreds of proteins that increase in abundance during aging, many of which also accumulate in the insoluble proteome. These datasets provide a valuable resource for prioritizing endogenous proteins that may contribute to age-associated proteostasis decline and serve as potential targets for pharmacological intervention. In the present study, we adopted a proteome-guided strategy to identify age-enriched proteins that could potentially be targeted by phytochemicals. We first analyzed the published age-dependent proteomic dataset of Walther et al. and identified 91 proteins whose abundance increased more than four-fold between day 1 and day 12 of adulthood. Human orthologs were identified, and the proteins were further evaluated for aggregation behavior and predicted aggregation propensity. Candidate proteins for molecular docking were prioritized based on several criteria, including functional characterization, conservation with human orthologs, substantial age-dependent accumulation, availability of structural information or closely related structural homologs, and biological relevance. Several proteins were selected using these criteria. However, in this study, we focused on glutamine-fructose-6-phosphate aminotransferase 2 (GFAT-2), a conserved enzyme that catalyzes the first and rate-limiting step of the hexosamine biosynthetic pathway (HBP). It is a nutrient-sensing branch of glucose metabolism that generates UDP-N-acetylglucosamine (UDP-GlcNAc), a key substrate for protein glycosylation and cellular signaling. The *C. elegans* genome encodes two glutamine:fructose-6-phosphate aminotransferase isoforms, GFAT-1 and GFAT-2, whereas the human genome contains two orthologous enzymes, GFPT1 (GFAT1) and GFPT2 (GFAT2), which catalyze the rate-limiting step of the HBP. The HBP has recently emerged as an important regulator of proteostasis and longevity. Activation of the HBP through gain-of-function mutations in its rate-limiting enzyme, glutamine:fructose-6-phosphate aminotransferase1 (GFAT-1), enhances endoplasmic reticulum (ER) stress resistance, reduce proteotoxicity, and extends lifespan in *C. elegans* (Denzel et al. 2014; Horn et al. 2020). Although the longevity-promoting effects of the HBP have been primarily attributed to GFAT-1, *C. elegans* GFAT-2 shares 88.3% amino acid sequence similarity with GFAT-1, suggesting that the two enzymes are likely to possess conserved structural and functional properties. GFAT-2 is homologous to human GFAT1 (GFPT1), mutations in which are associated with congenital myasthenic syndromes (Guergueltcheva et al. 2012; Kroef et al. 2022). Therefore, we selected GFAT-2 as a conserved target for phytochemical screening to evaluate whether compounds predicted to interact with this enzyme could mitigate age-associated protein aggregation.

We subsequently screened fifteen natural small molecules with reported therapeutic or longevity-promoting properties including caffeic acid, catechin, chlorogenic acid, p-coumaric acid, coumaroylquinic acid, curcumin, epicatechin, epigallocatechin gallate, feruloylquinic acid, hydroxybenzoic acid, kaempferol, quercetin, quinic acid, quinoline, and robigenin for their predicted interactions with GFAT-2. Among the compounds screened, quercetin exhibited the strongest predicted binding affinity toward both *C. elegans* GFAT-2 and its human ortholog, GFPT1. Quercetin is among the most extensively studied dietary flavonoids owing to its broad biological activities, including antioxidant, anti-inflammatory, and anti-aging effects (Klappan et al. 2012; Wang et al. 2021; Zhou et al. 2025). In *C. elegans*, quercetin has been reported to enhance stress tolerance and extend lifespan through conserved longevity pathways (Ayuda-Durán et al., 2019; Sugawara & Sakamoto 2020). Consequently, it has emerged as a promising candidate for mitigating age-associated proteotoxicity and protein aggregation (Yu & Lee 2020), although its effects on endogenous aggregation-prone proteins remain poorly understood. To determine whether the predicted interaction between quercetin and GFAT-2 translated into improved proteostasis in vivo, we evaluated the effect of quercetin on age-associated global protein aggregation by comparing soluble and insoluble protein fractions from young and aged worms following quercetin treatment. Although quercetin displayed favourable binding to GFAT, it did not significantly alter the accumulation of insoluble proteins during aging. These findings suggest that while quercetin may interact with specific aggregation-prone proteins, it does not measurably reduce the overall aggregation burden under the conditions tested. Nevertheless, our study establishes a systematic framework for integrating aging proteomics, structural biology, and molecular docking to identify endogenous proteins that accumulate during aging and evaluate phytochemicals as potential modulators of age-associated proteostasis. This approach provides a foundation for future studies investigating protein-specific aggregation mechanisms and the development of interventions targeting endogenous aggregation-prone proteins implicated in aging and age-related diseases.

## Methods

### Proteome data analysis

Age-associated changes in the *Caenorhabditis elegans* proteome were analyzed using the published quantitative proteomic datasets of Walther et al. (2015). Protein abundance, insoluble protein aggregation, and aggregation propensity datasets were examined. Proteins exhibiting a ≥4-fold increase in abundance at day 12 relative to day 1 were selected for further analyses. Age-dependent insoluble protein datasets were similarly compared between day 1 and day 12, while aggregation propensity was evaluated using the day 12 dataset. To facilitate cross-species comparison, *C. elegans* proteins were mapped to their corresponding human orthologs by retrieving Ensembl identifiers from WormBase, followed by conversion to official human gene symbols using Ensembl BioMart. Functional annotation and pathway enrichment analyses were performed using ShinyGO v0.85.1 (Ge et al., 2020). Statistical analyses and graphical visualizations were generated using R (v4.4.0) and associated packages.

### Protein sequence alignments

Protein sequences of *C. elegans* GFAT-2 and its human ortholog GFPT1 were retrieved from UniProt and aligned using CLUSTALW (Thompson et al., 1994) with default parameters. The resulting alignment file was visualized using ESPript 3.0 (Gouet et al., 1999) to identify conserved residues, and sequence similarity.

### Protein structure acquisition and validation

The predicted three-dimensional structure of *C. elegans* GFAT-2 was obtained from the AlphaFold Protein Structure Database (Abramson et al., 2024). The experimentally resolved human GFPT1 crystal structure was retrieved from the Protein Data Bank (Berman et al., 2000). Prior to structural analyses and molecular docking, both structures were prepared using BIOVIA Discovery Studio 2025 (Dassault Systèmes, 2025). Hydrogen atoms were added, incomplete side chains were corrected where necessary, and structures were subjected to energy minimization using the CHARMm force field to relieve steric clashes and optimize molecular geometry.

### Structural similarity analysis and validation

Structural conservation between *C. elegans* GFAT-2 and human GFPT1 was evaluated using UCSF Chimera v1.19 (Pettersen et al., 2021). Protein structures were superimposed by structural alignment, and root mean square deviation (RMSD) values were calculated to quantify structural similarity. Both pruned-pair RMSD, representing the conserved structural core, and all-pair RMSD, which includes flexible and divergent regions, were determined. Structural overlays were generated as ribbon diagrams for visualization.

The stereochemical quality of the predicted *C. elegans* GFAT-2 structure and the experimentally determined human GFPT1 structure was assessed using PROCHECK. Ramachandran plot analysis was performed to evaluate backbone dihedral angle distributions and overall structural quality. The majority of residues in both structures were located within the most favored regions, indicating that the models were suitable for subsequent molecular docking analyses.

### Selection of phytochemicals/natural small molecules

Fifteen phytochemicals with reported antioxidant, anti-aging, or stress-protective activities in *C. elegans* were selected for molecular docking (Table S4). Relevant literature was identified through PubMed searches using the query “*C. elegans* AND compound name.” Chemical structures, PubChem compound identifiers (CID), and SMILES strings were retrieved from PubChem (Kim et al., 2019) (Table S1). Pharmacokinetic and toxicity properties were predicted using SwissADME (Daina et al., 2017) and ADMETSAR 3.0 (Gu et al., 2024). The resulting datasets were compiled and compared based on physicochemical characteristics, absorption, distribution, metabolism, excretion, and toxicity parameters. Two-dimensional chemical structures used in the figures were generated using ChemDraw (PerkinElmer Informatics, 2003).

### Docking preparation and blind docking with selected phytochemicals/natural small molecules

Three-dimensional structures of all phytochemicals were downloaded from PubChem in Structure Data File (SDF) format. Ligands were imported into PyRx (Dallakyan & Olson, 2015), where they were subjected to geometry optimization and energy minimization using the Open Babel force field before conversion to PDBQT format for molecular docking.

Blind molecular docking was performed using PyRx (Dallakyan & Olson, 2015) with AutoDock Vina as the docking engine. Protein structures were prepared using UCSF Chimera v1.19 and BIOVIA Discovery Studio 2025 by removing non-essential molecules, adding hydrogen atoms, assigning charges, and performing energy minimization. Ligands were minimized within PyRx prior to docking. Docking calculations were performed using a grid box encompassing the entire protein structure to allow unbiased identification of potential binding sites. Binding affinities were recorded as predicted free energies of binding (kcal/mol). Docking scores were exported in CSV format for downstream analysis. Representative binding poses and molecular interactions, including hydrogen bonds, hydrophobic interactions, π-interactions, and electrostatic contacts, were visualized using BIOVIA Discovery Studio 2025.

### Maintenance of *C. elegans* strains

Wild-type *Caenorhabditis elegans* N2 worms were obtained from the Caenorhabditis Genetics Center (CGC, University of Minnesota, USA). Worms were maintained at 20°C on standard nematode growth medium (NGM) agar plates seeded with *Escherichia coli* OP50 following established protocols. Age-synchronized populations were generated by standard bleaching procedures.

### Quercetin treatment and collection of worms

Worms were exposed to 500 μM quercetin from the L1 larval stage until sample collection. Quercetin was dissolved in DMSO and added to the OP50 bacterial suspension before seeding onto NGM plates, resulting in a final DMSO concentration of 0.1%. Control animals received OP50 containing an equivalent concentration of DMSO. Worms were harvested at day 1 and day 10 of adulthood for biochemical analyses.

### Biochemical fractionation of soluble and insoluble proteome

Approximately 500 adult worms were collected and lysed in 450 μL of ice-cold lysis buffer containing 10 mM Tris-HCl (pH 7.5), 150 mM NaCl, 0.5 mM EDTA, 0.5% (v/v) NP-40, and a protease inhibitor cocktail. Worms were disrupted using a Bioruptor (Diagenode) sonication bath (30 min; 30 s ON/30 s OFF cycles; high power) at 4°C. Lysates were clarified by centrifugation at 1,000 × g for 2 min at 4°C to remove cuticle fragments and cellular debris.

Protein concentrations were determined using the Bradford assay and normalized prior to fractionation. Soluble and insoluble proteins were separated by ultracentrifugation (Eppendorf micro ultracentrifuge, CS150NX) at 120,000 × g for 20 min at 4°C. The supernatant was collected as the soluble protein fraction. The insoluble pellet was washed once with lysis buffer and solubilized in 0.5% SDS by incubation at 95°C for 10 min. Equal amounts of protein from soluble and insoluble fractions were resolved by SDS-PAGE and visualized by Coomassie Brilliant Blue staining.

## Results

### Identification of proteins that increase in abundance and aggregation during aging

To identify proteins that accumulate during aging, we analysed a published quantitative proteomic dataset comparing young (day 1) and aged (day 12) adult *C. elegans* (Walther et al. 2015). Proteins exhibiting a ≥4-fold increase in abundance during aging were selected for further analysis (Figure 1A). This analysis identified 220 proteins with increased abundance, of which 91 had identifiable human orthologs (Table 1). The selected proteins displayed varying degrees of age-dependent accumulation, with several exhibiting more than an eight-fold increase in abundance between day 1 and day 12 (Figure 1B). Visualization of the complete proteome using an MA plot confirmed that these proteins represented a distinct subset showing markedly increased abundance relative to the majority of quantified proteins (Figure 1C). Functional enrichment analysis revealed that these proteins were significantly associated with DNA replication and repair, metabolism and biosynthesis, longevity and stress response, signalling and structural processes, and protein degradation machinery (Figure 1D). To determine whether increased abundance was accompanied by protein aggregation, the selected proteins were compared with the age-associated insoluble proteome. Many of the identified proteins also accumulated in the insoluble fraction during aging (Figure 1E). Aggregation propensity data were available for 60 proteins, allowing simultaneous comparison of protein abundance, aggregation propensity, and aggregation load (Figure 1F). This analysis revealed substantial heterogeneity in the aggregation characteristics of age-associated proteins, with several proteins exhibiting both pronounced age-dependent accumulation and high aggregation load. Based on their abundance, aggregation behaviour, functional relevance, and conservation in humans, a few proteins were prioritized as candidate targets for subsequent computational analyses.

**Figure 1.**
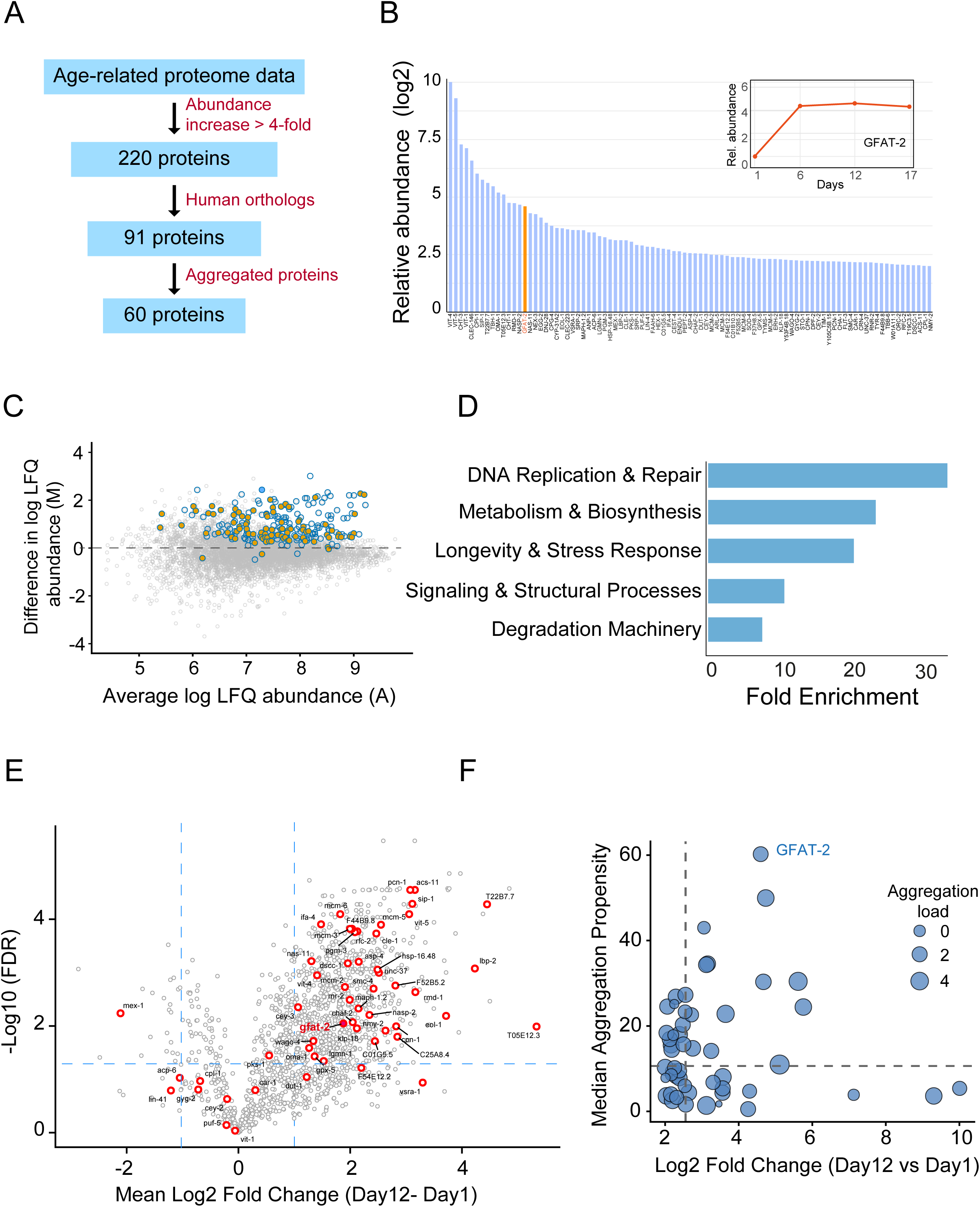
Proteome data analysis for identifying age-associated changes in protein abundance and aggregation and selecting candidate proteins for phytochemical screening. (A) Workflow used to identify candidate proteins from the published age-associated *C. elegans* proteome (Walther et al., 2015). Proteins showing a ≥4-fold increase in abundance between day 1 and day 12 adults were identified, mapped to human orthologs, and compared with the age-associated insoluble proteome to identify aggregation-prone candidates. (B) Relative abundance of proteins with human orthologs exhibiting a ≥4-fold increase in aged worms compared with young worms. Bars represent the log_2_ fold change in protein abundance (day 12/day 1) for each selected protein. (C) MA plot showing the distribution of age-associated proteins. The x-axis represents the average log_2_ protein abundance (A), and the y-axis represents the log_2_ abundance difference (M; day 12/day 1). Grey dots represent all quantified proteins, blue circles indicate proteins exhibiting a ≥4-fold increase during aging, and orange circles denote proteins with identified human orthologs. (D) Functional enrichment analysis of proteins with human orthologs. Significantly enriched functional categories are shown according to fold enrichment. (E) Volcano plot of proteins in the age-associated insoluble proteome. Grey circles represent all quantified proteins, whereas red circles indicate proteins with human orthologs. Dashed lines indicate the fold-change and significance thresholds. The x-axis represents the log_2_ fold change (day 12/day 1), and the y-axis represents the −log_10_ false discovery rate (FDR). (F) Relationship between protein abundance, aggregation propensity, and aggregation load. The x-axis represents the log_2_ fold change (day 12/day 1), the y-axis shows median aggregation propensity, bubble size corresponds to aggregation load, and bubble color indicates aggregation propensity. Representative proteins are labelled. Dashed lines denote the median values for abundance and aggregation propensity

**Table 1.**
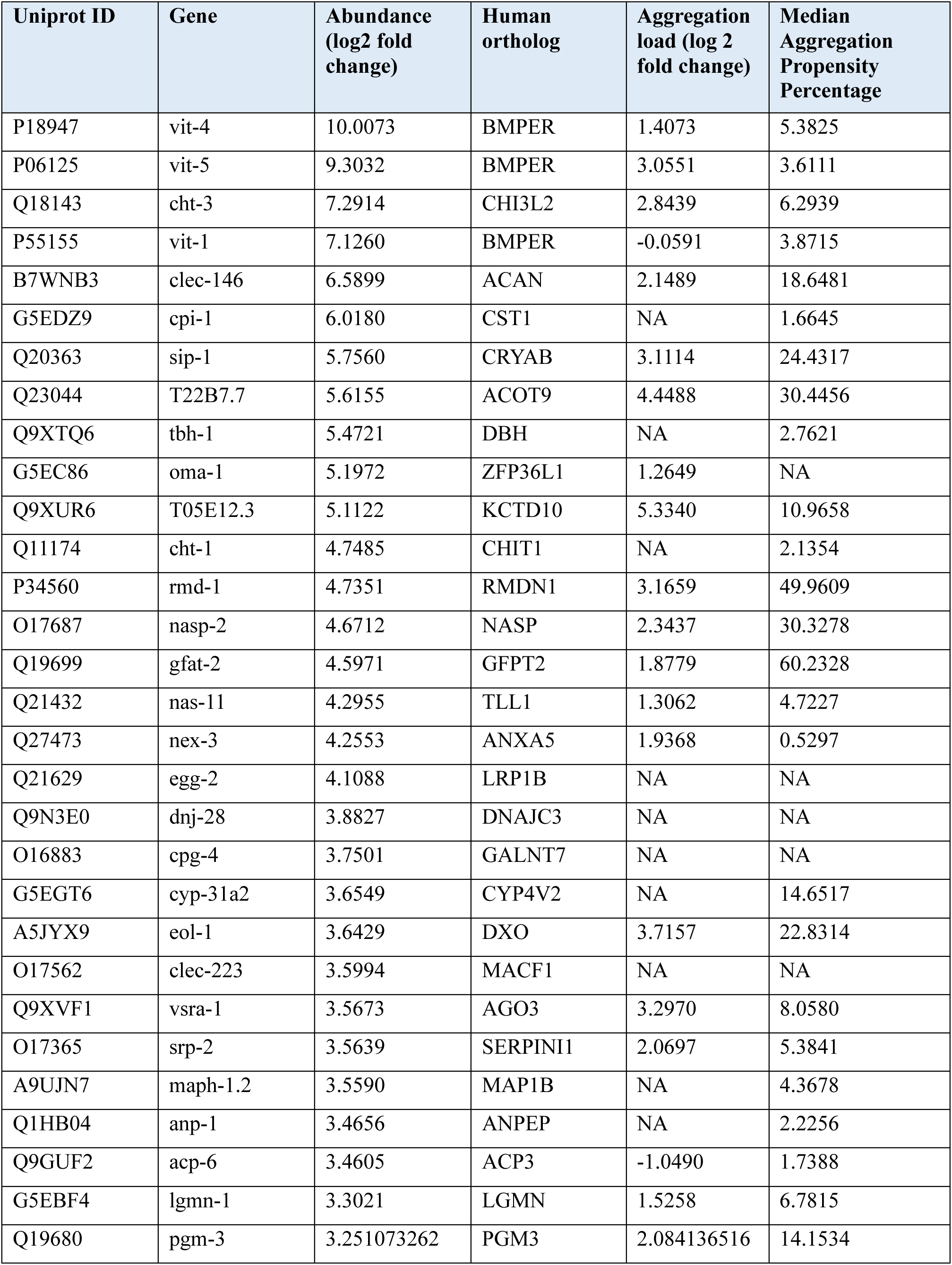
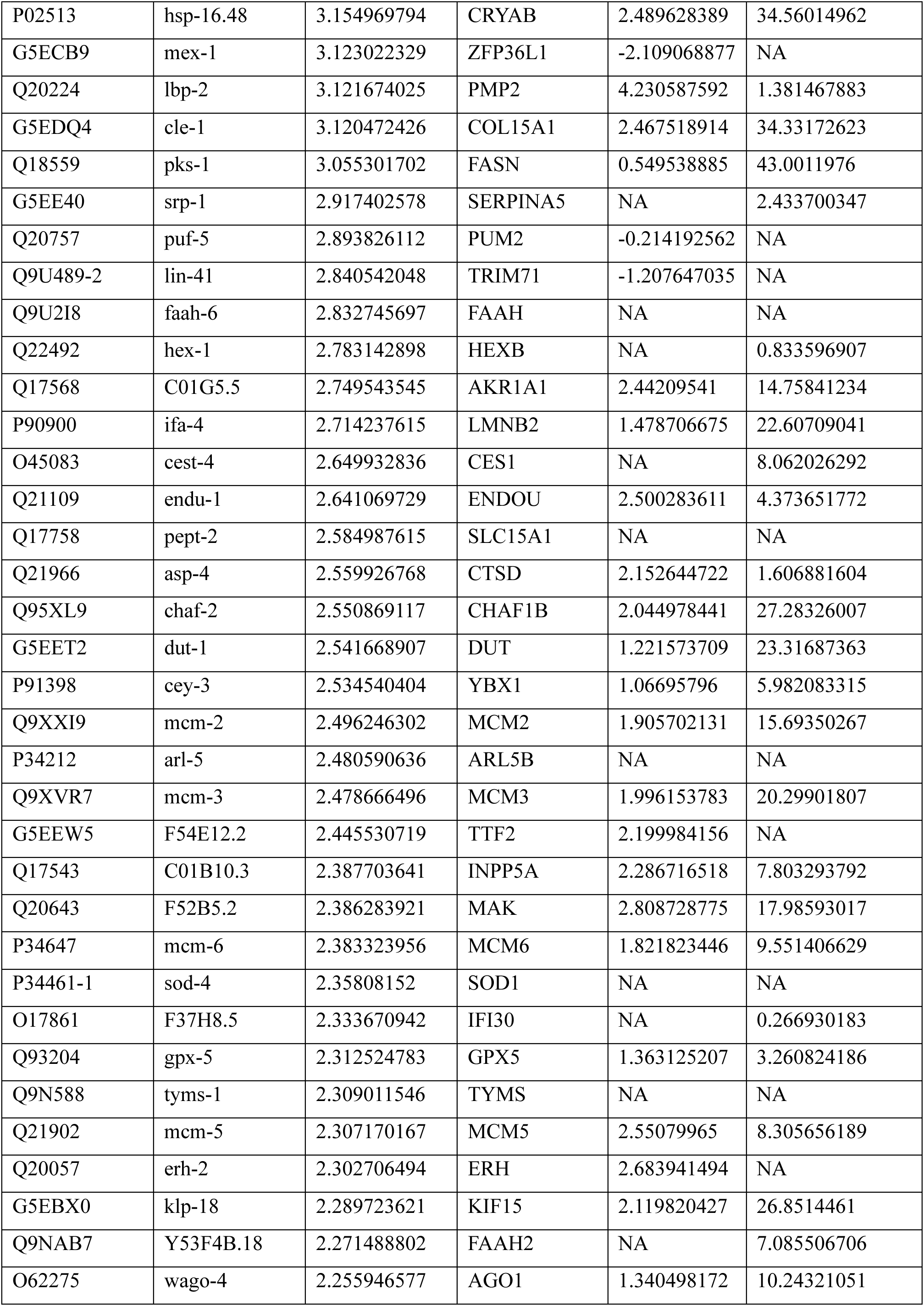
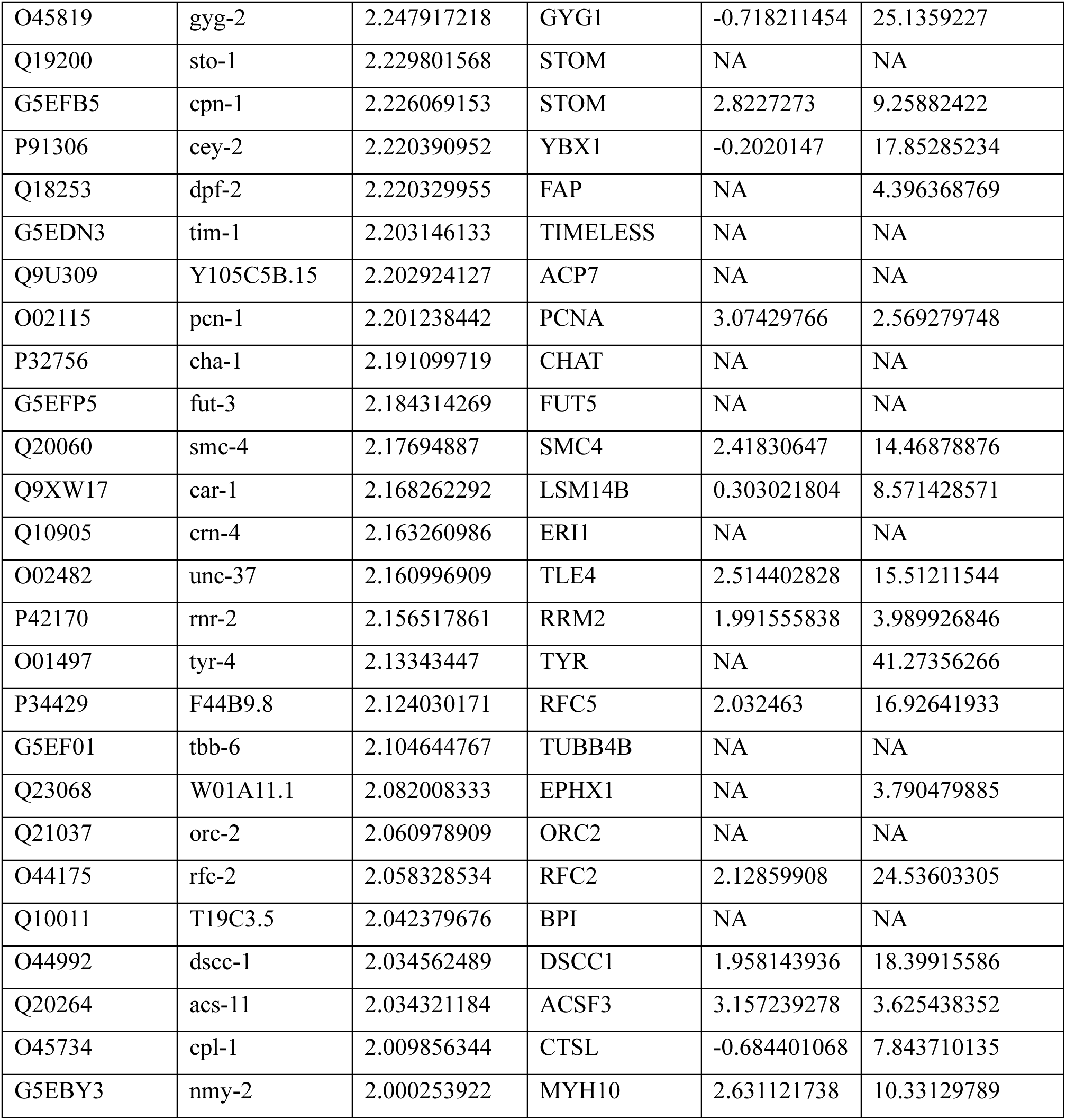
List of proteins that increase their abundance and aggregation.

| Uniprot ID | Gene | Abundance<br>(log2 fold<br>change) | Human<br>ortholog | Aggregation<br>load (log 2<br>fold change) | Median<br>Aggregation<br>Propensity<br>Percentage |
| --- | --- | --- | --- | --- | --- |
| P18947 | vit-4 | 10.0073 | BMPER | 1.4073 | 5.3825 |
| P06125 | vit-5 | 9.3032 | BMPER | 3.0551 | 3.6111 |
| Q18143 | cht-3 | 7.2914 | CHI3L2 | 2.8439 | 6.2939 |
| P55155 | vit-1 | 7.1260 | BMPER | -0.0591 | 3.8715 |
| B7WNB3 | clec-146 | 6.5899 | ACAN | 2.1489 | 18.6481 |
| G5EDZ9 | cpi-1 | 6.0180 | CST1 | NA | 1.6645 |
| Q20363 | sip-1 | 5.7560 | CRYAB | 3.1114 | 24.4317 |
| Q23044 | T22B7.7 | 5.6155 | ACOT9 | 4.4488 | 30.4456 |
| Q9XTQ6 | tbh-1 | 5.4721 | DBH | NA | 2.7621 |
| G5EC86 | oma-1 | 5.1972 | ZFP36L1 | 1.2649 | NA |
| Q9XUR6 | T05E12.3 | 5.1122 | KCTD10 | 5.3340 | 10.9658 |
| Q11174 | cht-1 | 4.7485 | CHIT1 | NA | 2.1354 |
| P34560 | rmd-1 | 4.7351 | RMDN1 | 3.1659 | 49.9609 |
| O17687 | nasp-2 | 4.6712 | NASP | 2.3437 | 30.3278 |
| Q19699 | gfat-2 | 4.5971 | GFPT2 | 1.8779 | 60.2328 |
| Q21432 | nas-11 | 4.2955 | TLL1 | 1.3062 | 4.7227 |
| Q27473 | nex-3 | 4.2553 | ANXA5 | 1.9368 | 0.5297 |
| Q21629 | egg-2 | 4.1088 | LRP1B | NA | NA |
| Q9N3E0 | dnj-28 | 3.8827 | DNAJC3 | NA | NA |
| O16883 | cpg-4 | 3.7501 | GALNT7 | NA | NA |
| G5EGT6 | cyp-31a2 | 3.6549 | CYP4V2 | NA | 14.6517 |
| A5JYX9 | eol-1 | 3.6429 | DXO | 3.7157 | 22.8314 |
| O17562 | clec-223 | 3.5994 | MACF1 | NA | NA |
| Q9XVF1 | vsra-1 | 3.5673 | AGO3 | 3.2970 | 8.0580 |
| O17365 | srp-2 | 3.5639 | SERPINI1 | 2.0697 | 5.3841 |
| A9UJN7 | maph-1.2 | 3.5590 | MAP1B | NA | 4.3678 |
| Q1HB04 | anp-1 | 3.4656 | ANPEP | NA | 2.2256 |
| Q9GUF2 | acp-6 | 3.4605 | ACP3 | -1.0490 | 1.7388 |
| G5EBF4 | lgmn-1 | 3.3021 | LGMN | 1.5258 | 6.7815 |
| Q19680 | pgm-3 | 3.251073262 | PGM3 | 2.084136516 | 14.1534 |
| P02513 | hsp-16.48 | 3.154969794 | CRYAB | 2.489628389 | 34.56014962 |
| G5ECB9 | mex-1 | 3.123022329 | ZFP36L1 | -2.109068877 | NA |
| Q20224 | lbp-2 | 3.121674025 | PMP2 | 4.230587592 | 1.381467883 |
| G5EDQ4 | cle-1 | 3.120472426 | COL15A1 | 2.467518914 | 34.33172623 |
| Q18559 | pks-1 | 3.055301702 | FASN | 0.549538885 | 43.0011976 |
| G5EE40 | srp-1 | 2.917402578 | SERPINA5 | NA | 2.433700347 |
| Q20757 | puf-5 | 2.893826112 | PUM2 | -0.214192562 | NA |
| Q9U489-2 | lin-41 | 2.840542048 | TRIM71 | -1.207647035 | NA |
| Q9U2I8 | faah-6 | 2.832745697 | FAAH | NA | NA |
| Q22492 | hex-1 | 2.783142898 | HEXB | NA | 0.833596907 |
| Q17568 | C01G5.5 | 2.749543545 | AKR1A1 | 2.44209541 | 14.75841234 |
| P90900 | ifa-4 | 2.714237615 | LMNB2 | 1.478706675 | 22.60709041 |
| O45083 | cest-4 | 2.649932836 | CES1 | NA | 8.062026292 |
| Q21109 | endu-1 | 2.641069729 | ENDOU | 2.500283611 | 4.373651772 |
| Q17758 | pept-2 | 2.584987615 | SLC15A1 | NA | NA |
| Q21966 | asp-4 | 2.559926768 | CTSD | 2.152644722 | 1.606881604 |
| Q95XL9 | chaf-2 | 2.550869117 | CHAF1B | 2.044978441 | 27.28326007 |
| G5EET2 | dut-1 | 2.541668907 | DUT | 1.221573709 | 23.31687363 |
| P91398 | cey-3 | 2.534540404 | YBX1 | 1.06695796 | 5.982083315 |
| Q9XXI9 | mcm-2 | 2.496246302 | MCM2 | 1.905702131 | 15.69350267 |
| P34212 | arl-5 | 2.480590636 | ARL5B | NA | NA |
| Q9XVR7 | mcm-3 | 2.478666496 | MCM3 | 1.996153783 | 20.29901807 |
| G5EEW5 | F54E12.2 | 2.445530719 | TTF2 | 2.199984156 | NA |
| Q17543 | C01B10.3 | 2.387703641 | INPP5A | 2.286716518 | 7.803293792 |
| Q20643 | F52B5.2 | 2.386283921 | MAK | 2.808728775 | 17.98593017 |
| P34647 | mcm-6 | 2.383323956 | MCM6 | 1.821823446 | 9.551406629 |
| P34461-1 | sod-4 | 2.35808152 | SOD1 | NA | NA |
| O17861 | F37H8.5 | 2.333670942 | IFI30 | NA | 0.266930183 |
| Q93204 | gpx-5 | 2.312524783 | GPX5 | 1.363125207 | 3.260824186 |
| Q9N588 | tyms-1 | 2.309011546 | TYMS | NA | NA |
| Q21902 | mcm-5 | 2.307170167 | MCM5 | 2.55079965 | 8.305656189 |
| Q20057 | erh-2 | 2.302706494 | ERH | 2.683941494 | NA |
| G5EBX0 | klp-18 | 2.289723621 | KIF15 | 2.119820427 | 26.8514461 |
| Q9NAB7 | Y53F4B.18 | 2.271488802 | FAAH2 | NA | 7.085506706 |
| O62275 | wago-4 | 2.255946577 | AGO1 | 1.340498172 | 10.24321051 |
| O45819 | gyg-2 | 2.247917218 | GYG1 | -0.718211454 | 25.1359227 |
| Q19200 | sto-1 | 2.229801568 | STOM | NA | NA |
| G5EFB5 | cpn-1 | 2.226069153 | STOM | 2.8227273 | 9.25882422 |
| P91306 | cey-2 | 2.220390952 | YBX1 | -0.2020147 | 17.85285234 |
| Q18253 | dpf-2 | 2.220329955 | FAP | NA | 4.396368769 |
| G5EDN3 | tim-1 | 2.203146133 | TIMELESS | NA | NA |
| Q9U309 | Y105C5B.15 | 2.202924127 | ACP7 | NA | NA |
| O02115 | pcn-1 | 2.201238442 | PCNA | 3.07429766 | 2.569279748 |
| P32756 | cha-1 | 2.191099719 | CHAT | NA | NA |
| G5EFP5 | fut-3 | 2.184314269 | FUT5 | NA | NA |
| Q20060 | smc-4 | 2.17694887 | SMC4 | 2.41830647 | 14.46878876 |
| Q9XW17 | car-1 | 2.168262292 | LSM14B | 0.303021804 | 8.571428571 |
| Q10905 | crn-4 | 2.163260986 | ERI1 | NA | NA |
| O02482 | unc-37 | 2.160996909 | TLE4 | 2.514402828 | 15.51211544 |
| P42170 | rnr-2 | 2.156517861 | RRM2 | 1.991555838 | 3.989926846 |
| O01497 | tyr-4 | 2.13343447 | TYR | NA | 41.27356266 |
| P34429 | F44B9.8 | 2.124030171 | RFC5 | 2.032463 | 16.92641933 |
| G5EF01 | tbb-6 | 2.104644767 | TUBB4B | NA | NA |
| Q23068 | W01A11.1 | 2.082008333 | EPHX1 | NA | 3.790479885 |
| Q21037 | orc-2 | 2.060978909 | ORC2 | NA | NA |
| O44175 | rfc-2 | 2.058328534 | RFC2 | 2.12859908 | 24.53603305 |
| Q10011 | T19C3.5 | 2.042379676 | BPI | NA | NA |
| O44992 | dsc-1 | 2.034562489 | DSCC1 | 1.958143936 | 18.39915586 |
| Q20264 | acs-11 | 2.034321184 | ACSF3 | 3.157239278 | 3.625438352 |
| O45734 | cpl-1 | 2.009856344 | CTSL | -0.684401068 | 7.843710135 |
| G5EBY3 | nmy-2 | 2.000253922 | MYH10 | 2.631121738 | 10.33129789 |

**Table 2.**
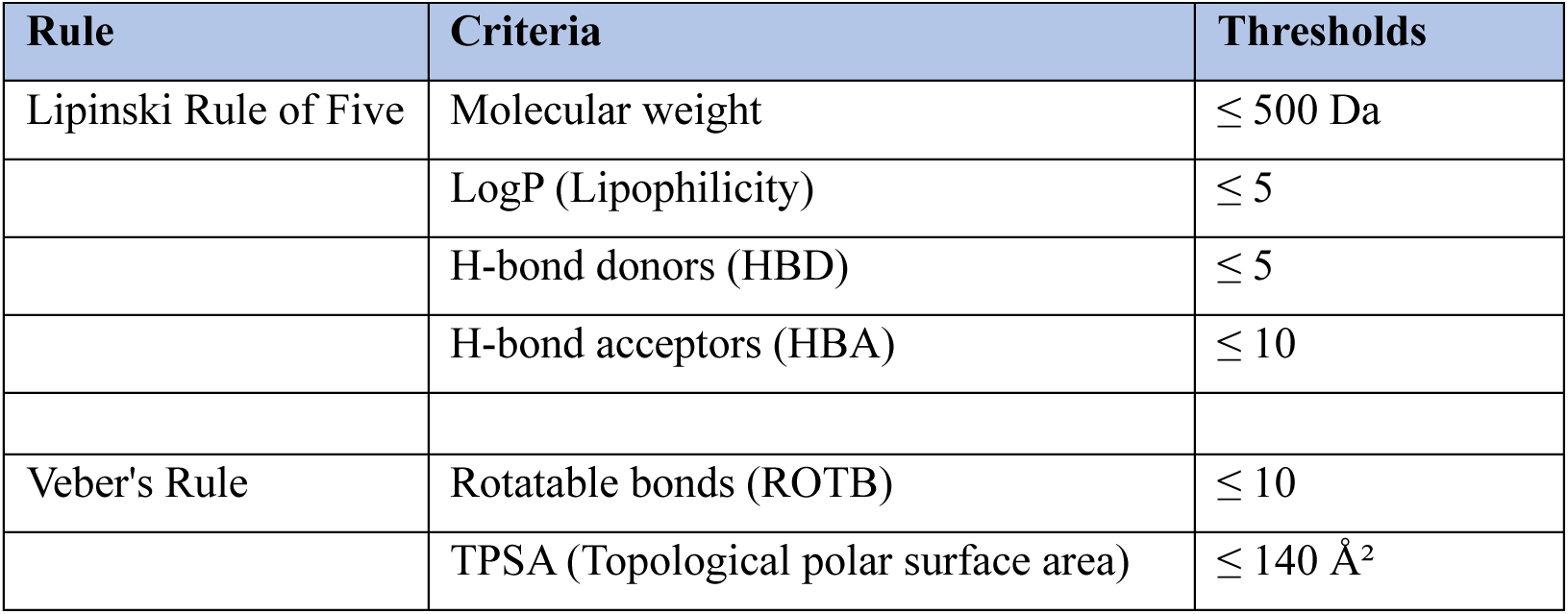
Thresholds defined by Lipinski’s and Veber’s rule.

**Table 3.** Physiochemical filter defining oral drug likeliness (Lipinski and Veber’s Rule)

| Molecule | MW | iLOGP | HBD | HBA | ROTB | TPSA | Lipinski #violations | Veber #violations |
| --- | --- | --- | --- | --- | --- | --- | --- | --- |
| Caffeic acid | 180.16 | 0.97 | 3 | 4 | 2 | 77.76 | 0 | 0 |
| Catechin | 290.27 | 1.36 | 5 | 6 | 1 | 110.38 | 0 | 0 |
| Chlorogenic acid | 354.31 | 0.87 | 6 | 9 | 5 | 164.75 | 1 | 1 |
| Coumaric acid | 164.16 | 0.95 | 2 | 3 | 2 | 57.53 | 0 | 0 |
| Coumaroylquinic acid | 338.31 | 0.83 | 5 | 8 | 5 | 144.52 | 0 | 1 |
| Curcumin | 368.38 | 3.27 | 2 | 6 | 8 | 93.06 | 0 | 0 |
| Epicatechin | 290.27 | 1.47 | 5 | 6 | 1 | 110.38 | 0 | 0 |
| Epigallocatechingallate | 458.37 | 1.53 | 8 | 11 | 4 | 197.37 | 2 | 1 |
| Feruloyl quinic acid | 368.34 | 0.92 | 5 | 9 | 6 | 153.75 | 0 | 1 |
| Hydrobenzoic acid | 138.12 | 0.86 | 2 | 3 | 1 | 57.53 | 0 | 0 |
| Kaempferol | 418.35 | 1.85 | 6 | 10 | 3 | 170.05 | 1 | 1 |
| Quercetin | 448.38 | 1.6 | 7 | 11 | 3 | 190.28 | 2 | 1 |
| Quinic acid | 192.17 | 0.37 | 5 | 6 | 1 | 118.22 | 0 | 0 |
| Quinoline | 159.19 | 1.1 | 2 | 1 | 0 | 64.93 | 0 | 0 |
| Robigenin | 286.24 | 1.7 | 4 | 6 | 1 | 111.13 | 0 | 0 |

Abbreviations:
MW - Molecular Weight, iLOGP - Partition coefficient between octanol and water, TPSA - Topological Polar Surface Area, ROTB - Rotatable bonds, HBD - Hydrogen Bond Donor, HBA - Hydrogen Bond Acceptor.

**Table 4.** Pharmacokinetic parameter related to distribution and metabolism.

| Molecule | Pgp substrate | BBB permeant | GI absorption | CYP3A4 inhibitor | CYP1A2 inhibitor | CYP2D6 inhibitor |
| --- | --- | --- | --- | --- | --- | --- |
| Caffeic acid | No | No | High | No | No | No |
| Catechin | Yes | No | High | No | No | No |
| Chlorogenic acid | No | No | Low | No | No | No |
| Coumaric acid | No | Yes | High | No | No | No |
| Coumaroylquinic acid | No | No | Low | No | No | No |
| Curcumin | No | No | High | Yes | No | No |
| Epicatechin | Yes | No | High | No | No | No |
| Epigallocatechingallate | No | No | Low | No | No | No |
| Feruloyl quinic acid | No | No | Low | No | No | No |
| Hydrobenzoic acid | No | Yes | High | No | No | No |
| Kaempferol | No | No | Low | No | No | No |
| Quercetin | No | No | Low | No | No | No |
| Quinic acid | No | No | Low | No | No | No |
| Quinoline | No | Yes | High | No | Yes | No |
| Robigenin | No | No | High | Yes | Yes | Yes |

**Table 5.** Interaction type between protein candidate and selected phytochemical.

| Protein | Type | Interactors | Amino acids | Bond type |
| --- | --- | --- | --- | --- |
| GFAT-2 | *C. elegans* | Quercetin (-9.7kcal/mol) | Gln 450, Ser 502, Val 707, Lys586, Cys402, Thr404, Ser 449 | Conventional hydrogen bond, Carbon hydrogen bond, Unfavourable donor-donor, Pi-Sigma, Pi-alkyl |
| GFPT1 | Humans | Quercetin (-8.6kcal/mol) | Tyr549, Tyr378, Tyr549, Gly569, Glu568, Leu553, Leu557, Lys560 | Conventional hydrogen bond, Pi-anion, Unfavourable donor-donor, Pi-alkyl |

### Selection of GFAT protein as a conserved target for phytochemical screening

To identify a suitable target for phytochemical screening, proteins were prioritized based on biological relevance, structural availability, conservation with human orthologs, and suitability for molecular docking. Candidate proteins were required to (i) exhibit at least a four-fold increase in abundance during aging, (ii) possess a human ortholog, (iii) have an experimentally determined structure or a reliable structural model, and (iv) perform an important biological function. Following this pre-screening, several proteins were selected (Table 1). In this study, we selected, glutamine:fructose-6-phosphate aminotransferase-2 (GFAT-2) for further investigation. The *C. elegans* genome encodes two glutamine:fructose-6-phosphate aminotransferase isoforms, GFAT-1 and GFAT-2, whereas the human genome contains two orthologous enzymes, GFPT1 (GFAT1) and GFPT2 (GFAT2), which catalyse the rate-limiting step of the hexosamine biosynthetic pathway. We also found GFAT-1 aggrgegation during aging. Two other independent studies also confirmed age dependent aggregation of GFAT-2 (David et al. 2010; Reis-Rodrigues et al. 2012) and GFAT-1 (David et al. 2010). Both, GFAT-1 and GFAT-2 catalyse the rate-limiting step of the hexosamine biosynthetic pathway and are homologous to human GFPT1/2, highlighting thier biological relevance and translational potential.

Sequence alignment demonstrated extensive conservation between *C. elegans* GFAT-2 and human GFPT1 (Figure 2A). Structural superimposition of the AlphaFold-predicted *C. elegans* GFAT-2 model with the human crystal structure revealed a high degree of structural similarity, particularly within the conserved catalytic core (Figure 2B). Ramachandran plot analysis confirmed the stereochemical quality of the predicted *C. elegans* structure, supporting its suitability for molecular docking (Figure 2C).

**Figure 2.**
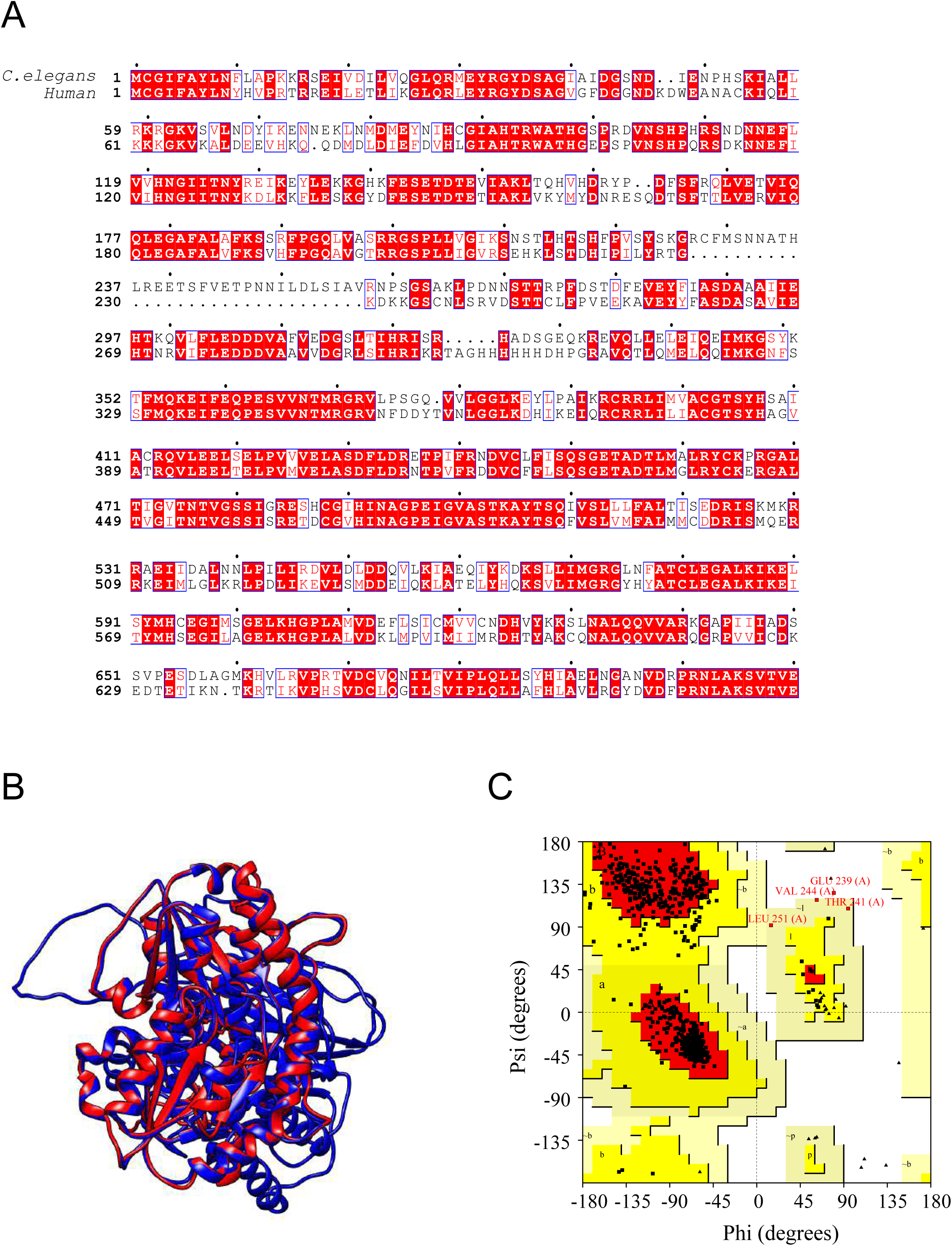
Sequence and structural comparison of *C. elegans* GFAT-2 and human GFPT1. (A) Sequence conservation between *C. elegans* GFAT-2 and human GFPT1. Multiple sequence alignment was generated using ClustalW and visualized with ESPript. Conserved residues are highlighted in red, demonstrating the high sequence conservation between the two orthologs. (B) Structural superimposition of *C. elegans* GFAT-2 and human GFPT1. The AlphaFold-predicted *C. elegans* GFAT-2 structure was superimposed onto the human GFPT1 crystal structure, demonstrating a high degree of structural conservation (RMSD = 0.523 Å after refinement). (C) Ramachandran plot validation of the predicted *C. elegans* GFAT-2 structure. PROCHECK analysis showed that 90.3% of residues were located in the most favoured regions and 99.3% in favoured or allowed regions, supporting the quality of the predicted structure for molecular docking.

### Physicochemical characterization of selected phytochemicals

Fifteen phytochemicals previously reported to possess antioxidant, stress-protective, or lifespan-promoting activities in *C. elegans* were selected for molecular docking (Table S2). Their physicochemical properties were evaluated according to Lipinski’s and Veber’s drug-likeness criteria. Radar plot analysis demonstrated considerable structural diversity among the selected compounds (Figure 3A). Most phytochemicals complied with the recommended ranges for molecular weight, lipophilicity, hydrogen-bond donor and acceptor numbers, rotatable bonds, and topological polar surface area. However, several highly hydroxylated polyphenols exhibited relatively high hydrogen-bonding capacity and topological polar surface area, reflecting their greater polarity.

**Figure 3.**
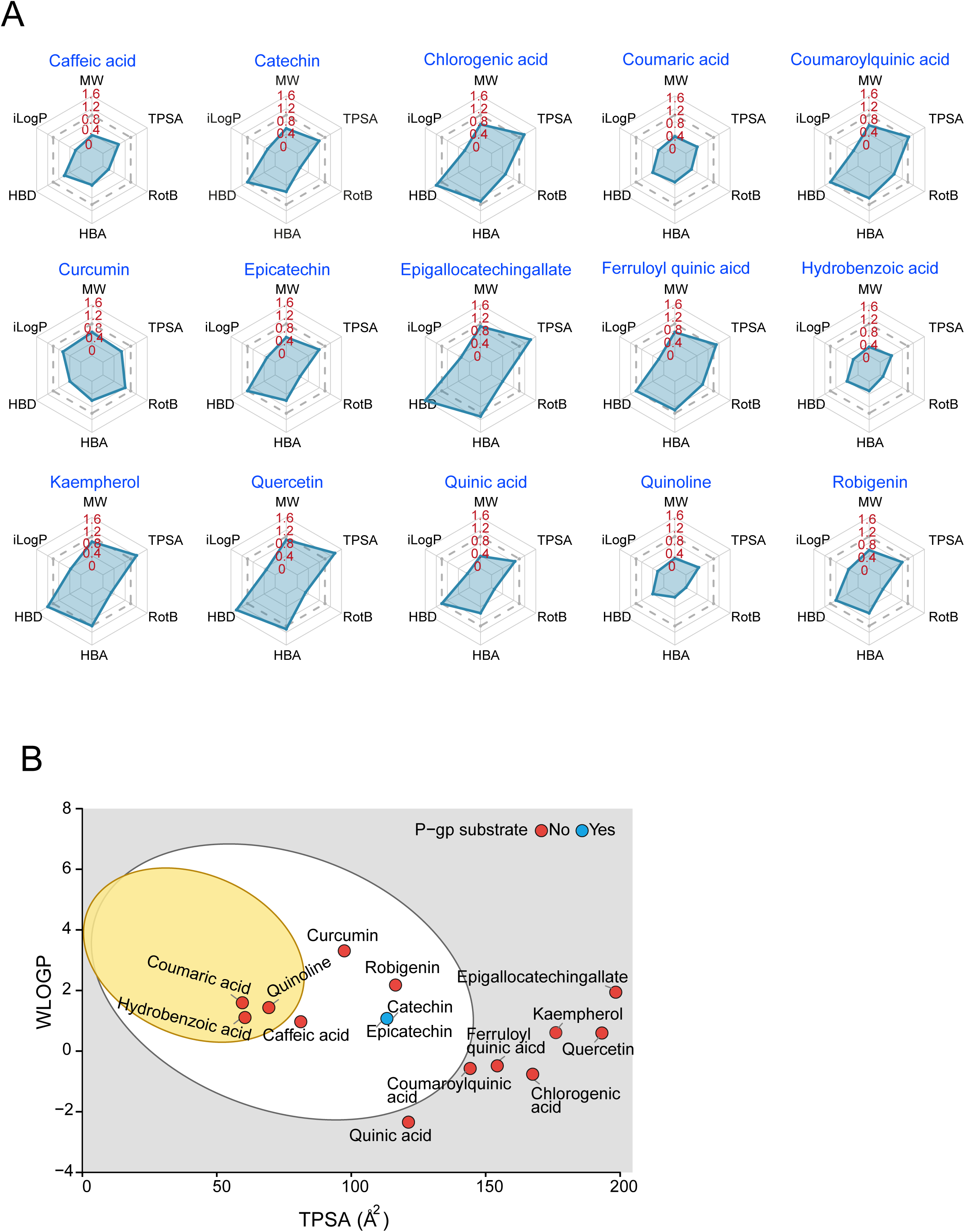
Physicochemical properties and predicted pharmacokinetic characteristics of the selected phytochemicals. (A) Radar plots showing the normalized physicochemical properties of the fifteen phytochemicals. The parameters include molecular weight (MW), lipophilicity (iLogP), hydrogen bond donors (HBD), hydrogen bond acceptors (HBA), rotatable bonds (RotB), and topological polar surface area (TPSA). The dashed polygon represents the recommended Lipinski/Veber drug-likeness thresholds, with values extending beyond the boundary indicating violations of these criteria. (B) BOILED-Egg model predicting passive gastrointestinal absorption and blood–brain barrier (BBB) permeability of the selected phytochemicals. The white region indicates compounds predicted to exhibit high gastrointestinal absorption, whereas the yellow region represents compounds predicted to penetrate the BBB. Symbols are coloured according to their predicted P-glycoprotein (P-gp) substrate status.

Predicted pharmacokinetic properties were further assessed using the BOILED-Egg model (Figure 3B). Nine phytochemicals were predicted to possess high gastrointestinal absorption, whereas epigallocatechin gallate, quercetin, kaempferol, chlorogenic acid, feruloyl quinic acid, and coumaroylquinic acid were predicted to have lower passive intestinal absorption because of their relatively high polarity. Catechin was the only compound predicted to be a P-glycoprotein substrate.

### Comparative molecular docking identifies quercetin as the highest-affinity ligand

The fifteen phytochemicals were docked against *C. elegans* GFAT-2 and its human ortholog GFPT1 to identify compounds with strong and conserved binding affinity. Among all compounds tested, quercetin exhibited the strongest predicted binding affinity for both proteins, with docking scores of −9.7 kcal/mol for *C. elegans* GFAT-2 and −8.6 kcal/mol for human GFPT1 (Figures 4A and 4B). Kaempferol, robigenin, catechin, epicatechin, and epigallocatechin gallate also displayed relatively strong binding affinities (Suppl Figure 1), whereas quinoline, chlorogenic acid, hydroxybenzoic acid, and feruloyl quinic acid showed comparatively weaker interactions.

**Figure 4.**
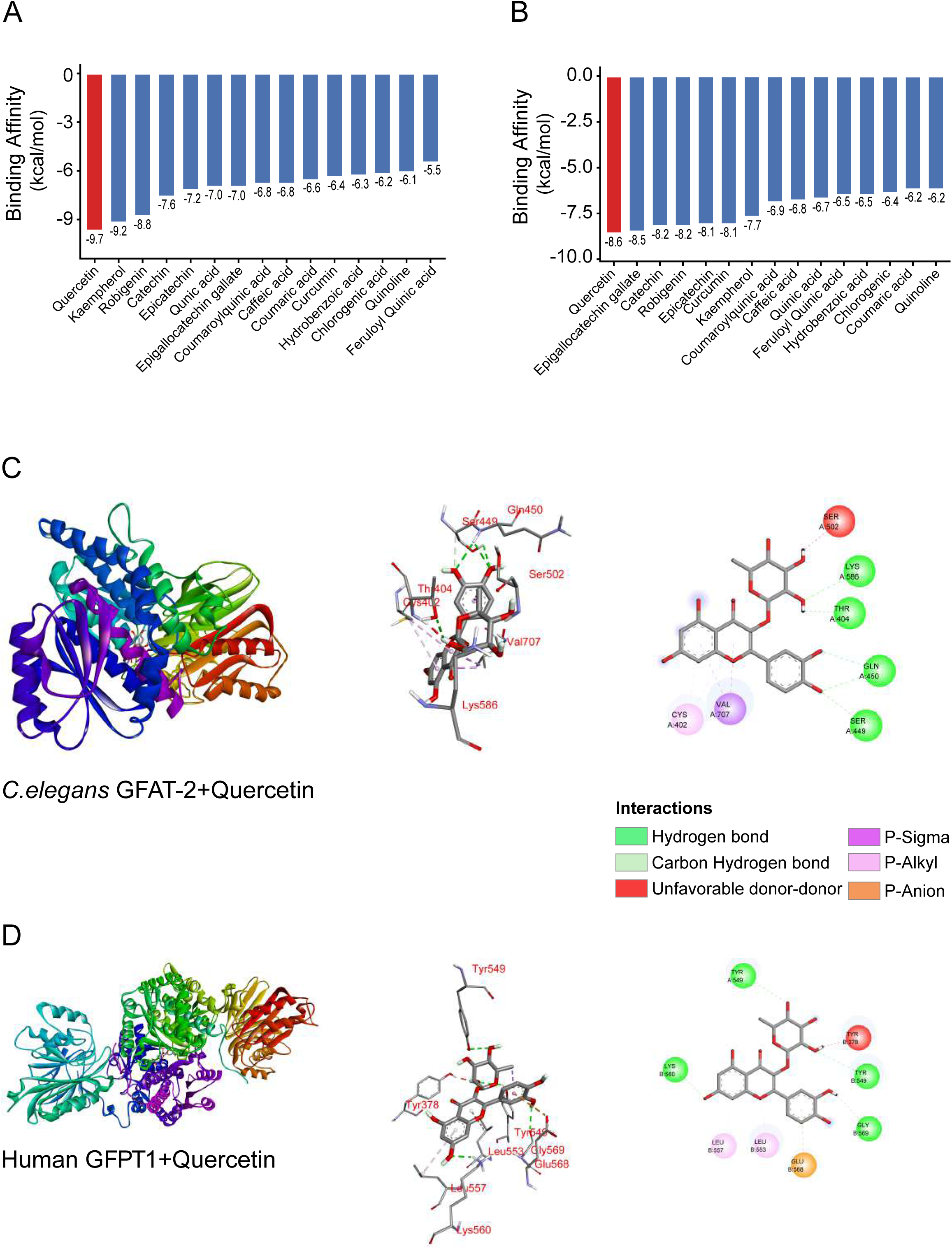
Molecular docking analysis of phytochemicals with *C. elegans* GFAT-2 and human GFPT1. (A) Predicted binding affinities of fifteen phytochemicals docked against *C. elegans* GFAT-2. Compounds are ranked according to binding energy (kcal/mol), with more negative values indicating stronger predicted binding. (B) Predicted binding affinities of the same phytochemicals against human GFPT1. Quercetin exhibited the strongest predicted binding affinity for both orthologs. (C) Predicted binding mode of quercetin with *C. elegans* GFAT-2. Left, overall protein structure showing the docked ligand; middle, enlarged view of the binding pocket; right, two-dimensional interaction diagram illustrating hydrogen bonds, carbon-hydrogen bonds, π-interactions, and other residue–ligand contacts. (D) Predicted binding mode of quercetin with human GFPT1 presented as described for panel C.

Because quercetin consistently showed the highest binding affinity for both orthologs, it was selected for detailed interaction analysis. Docking predicted that quercetin binds within a conserved binding pocket in both proteins (Figures 4C and 4D). In *C. elegans* GFAT-2, the complex was stabilized by multiple hydrogen bonds together with carbon-hydrogen, π–σ, π–alkyl, and electrostatic interactions involving residues within the binding pocket (Figure 4C). A similar interaction pattern was observed for human GFPT1, indicating that the predicted binding mode is highly conserved between the two orthologs. (Figure 4D).

### Effect of quercetin on global protein insolubility during aging

To determine whether the strong predicted interaction between quercetin and GFAT-2 translated into altered proteostasis in vivo, age-synchronized worms were exposed to 500 μM quercetin from the L1 stage, and soluble and insoluble protein fractions were isolated from day 1 and day 10 adults. Representative SDS-PAGE analysis showed the expected age-dependent increase in protein insolubility but did not reveal obvious differences between untreated, vehicle-treated (DMSO), and quercetin-treated worms (Figure 5A). Quantification of the insoluble protein fractions confirmed that quercetin treatment did not significantly alter the amount of aggregated protein at either day 1 or day 10, irrespective of whether samples were analyzed in the presence or absence of SDS (Figures 5B and 5C). Thus, despite exhibiting the strongest predicted binding affinity toward GFAT-2 in silico, quercetin did not measurably alter global age-associated protein insolubility under the experimental conditions used in this study. Although quercetin did not reduce global protein insolubility, it may modulate inappropriate protein-protein interactions without altering the overall aggregation burden. These findings highlight that favourable in silico docking predictions require experimental validation.

**Figure 5.**
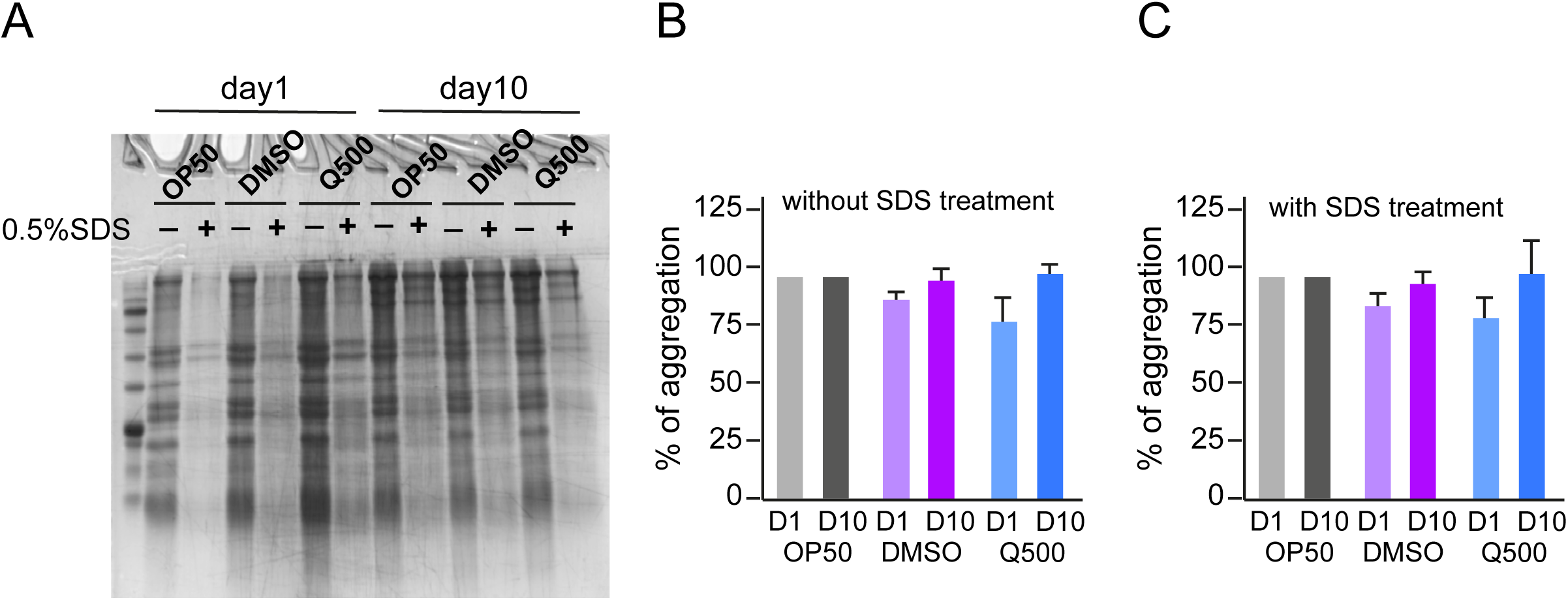
Quercetin does not alter global protein insolubility during aging in *C. elegans*. (A) Representative SDS-PAGE analysis of soluble and insoluble protein fractions isolated from day 1 and day 10 adult worms maintained on OP50, vehicle control (0.1% DMSO), or 500 μM quercetin (Q500). Protein fractions were analysed in the presence or absence of SDS treatment. (B) Quantification of protein aggregation in samples analysed without SDS treatment. (C) Quantification of protein aggregation following SDS treatment. Protein aggregation increased with age; however, no significant differences were observed between quercetin-treated worms and the corresponding control groups.

## Discussion

Loss of proteostasis is a hallmark of aging and is characterized by the progressive accumulation of misfolded and insoluble proteins due to declining protein quality control mechanisms. Although numerous studies have catalogued proteins that aggregate during aging, comparatively few have explored whether endogenous age-associated aggregation-prone proteins can be targeted by small molecules. In this study, we integrated published proteomic datasets, structural bioinformatics, molecular docking, and experimental validation to identify candidate proteins that accumulate during aging and evaluate phytochemicals as potential modulators of their aggregation. By analyzing the age-dependent proteome of *C. elegans*, we identified 91 proteins whose abundance increased more than four-fold during aging. A substantial proportion of these proteins also accumulated in the insoluble proteome, supporting previous observations that increased protein abundance contributes to age-associated aggregation.

Among the identified proteins, GFAT-2 was selected for further investigation because of its age-dependent accumulation, conservation with the human ortholog GFPT1, and availability of high-confidence structural models. GFAT catalyzes the rate-limiting step of the hexosamine biosynthetic pathway, which regulates nutrient sensing and protein glycosylation. Increasing evidence suggests that modulation of this pathway influences stress resistance, metabolism, and longevity, making GFAT-2 an attractive target fo r investigating phytochemical interactions. Screening of fifteen phytochemicals identified quercetin as the compound with the strongest predicted binding affinity toward both *C. elegans* and human GFAT proteins. Other natural molecules such as Kaempferol, robigenin, catechin, epicatechin, and epigallocatechin gallate also showed strong binding affinities. The conserved binding mode observed in both proteins suggests that the interaction is evolutionarily preserved and supports the use of *C. elegans* as a model for studying conserved protein–ligand interactions.

Quercetin has previously been reported to improve stress resistance and lifespan in several experimental systems. However, biochemical analyses demonstrated that quercetin did not significantly alter global protein insolubility during aging. One possible explanation is that quercetin interacts with multiple aggregation-prone proteins rather than a single target. By binding to exposed hydrophobic or aggregation-prone regions of diverse proteins, quercetin may reduce inappropriate intermolecular interactions that promote proteotoxicity without necessarily decreasing the total amount of insoluble protein. Thus, quercetin may improve the quality or stability of protein interactions rather than altering the overall aggregation burden.

This study also demonstrates the importance of experimentally validating computational predictions. While molecular docking provides a valuable approach for prioritizing candidate compounds, biochemical validation remains essential before biological significance can be established. Future studies should directly examine the aggregation, localization, and enzymatic activity of GFAT-2 in quercetin-treated animals using protein-specific approaches. Additional biophysical techniques, such as thermal shift assays or surface plasmon resonance, together with molecular dynamics simulations, would further validate the predicted interaction.

The present study has several limitations. Only one candidate protein was investigated experimentally, and aggregation was assessed at the global proteome level rather than for individual proteins. Furthermore, although quercetin showed no effect on total protein insolubility, its influence on specific aggregation-prone proteins or other proteostasis pathways cannot be excluded. Despite these limitations, this work establishes a proteome-guided framework for identifying endogenous proteins that accumulate during aging and evaluating phytochemicals that may target them. Rather than focusing solely on disease-associated proteins, this approach identifies physiologically relevant proteins that naturally aggregate during aging and therefore may represent novel therapeutic targets. Importantly, our findings emphasize that computational predictions should be complemented by rigorous experimental validation.

## Acknowledgements

This work was supported by funding from Department of Biotechnology, Government of India, as DBT-Ramalingaswami Re-entry grant (BT/RLF/Re-entry/31/2018 to P.K). The authors gratefully acknowledge the support of the DST, Government of India, for providing infrastructure facilities through the FIST program (Grant No. SR/FST/LS-II/2022/950).

---

**Table S1.**
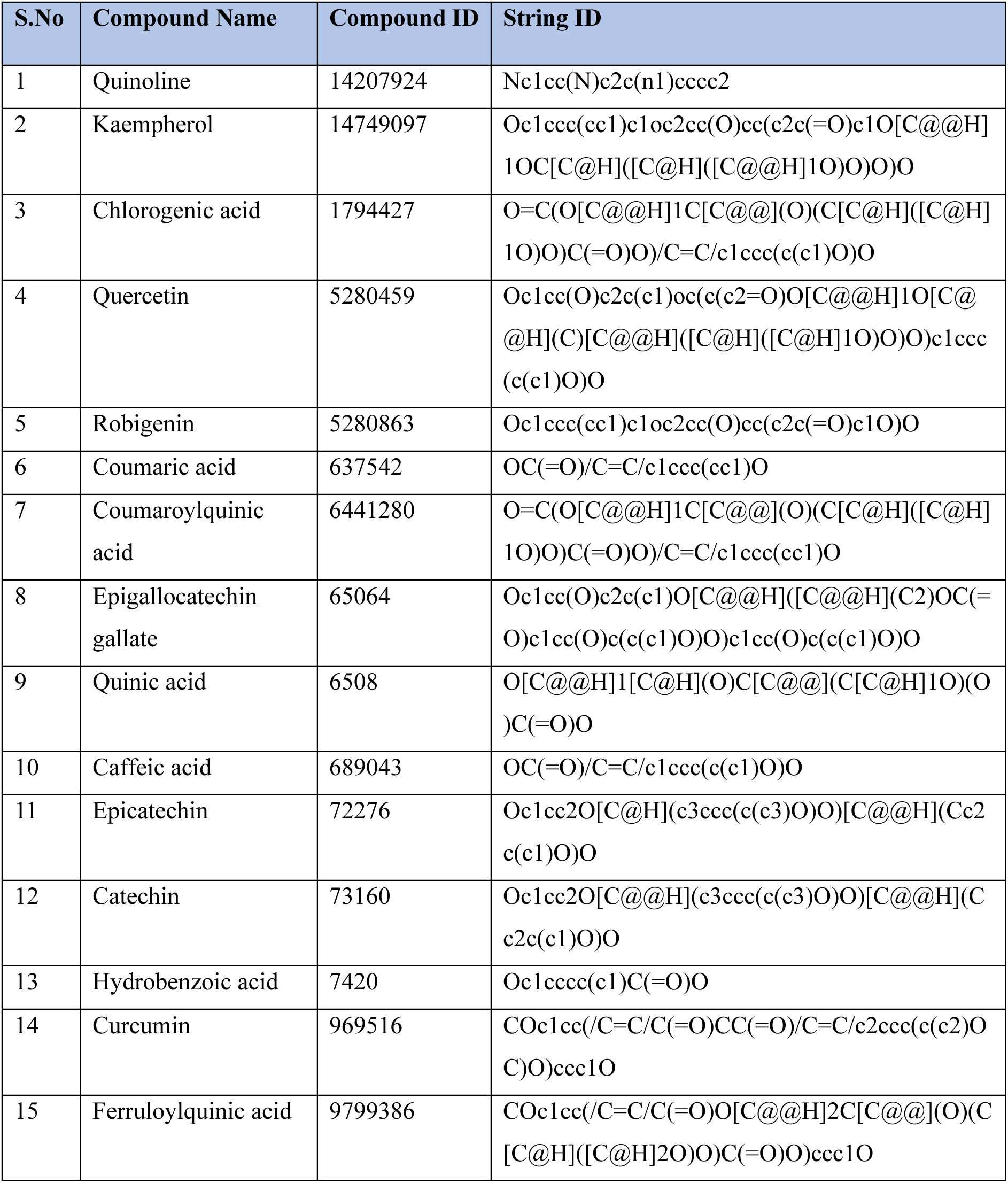
STRING ID for phytochemicals.

**Table S2.**
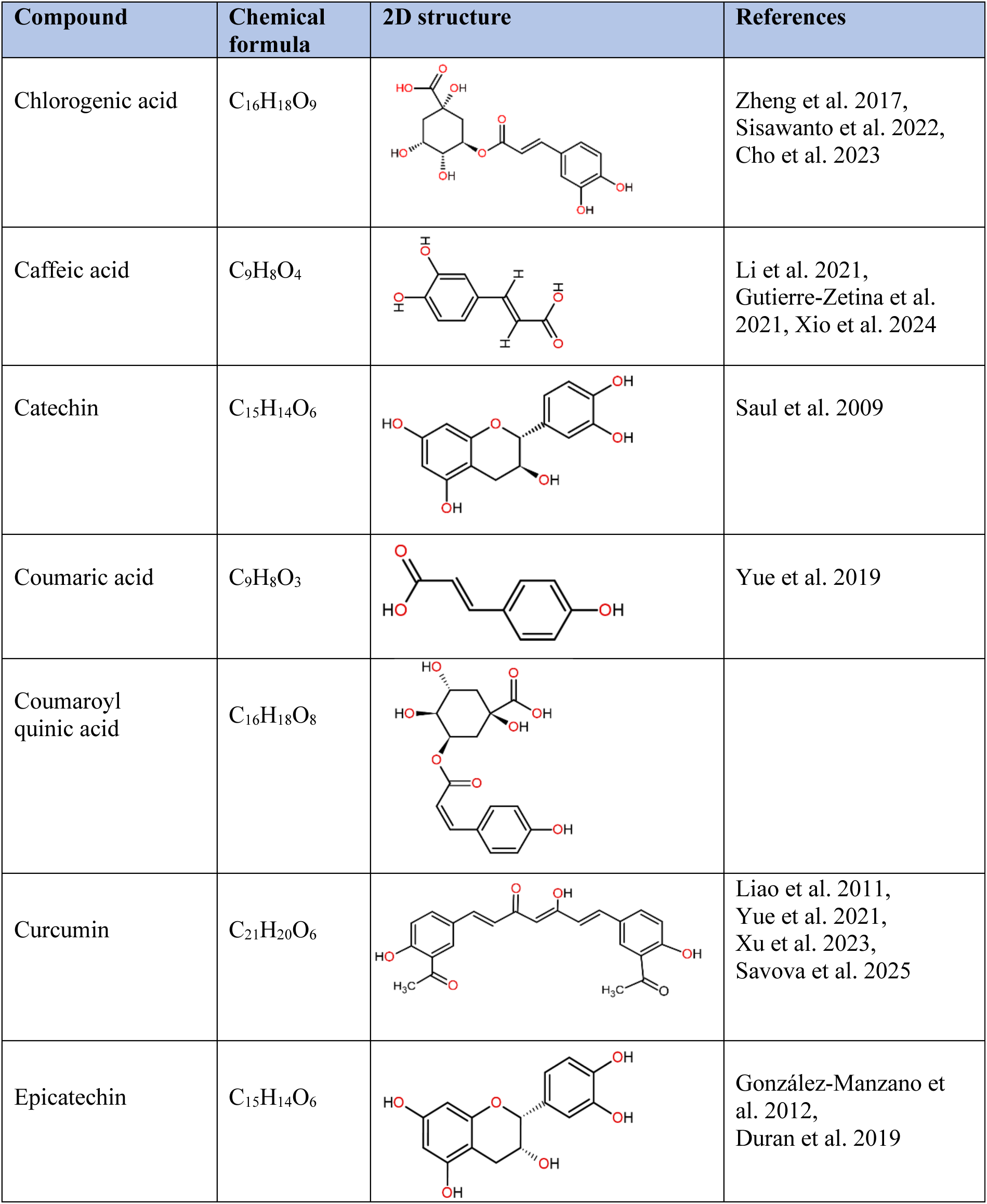
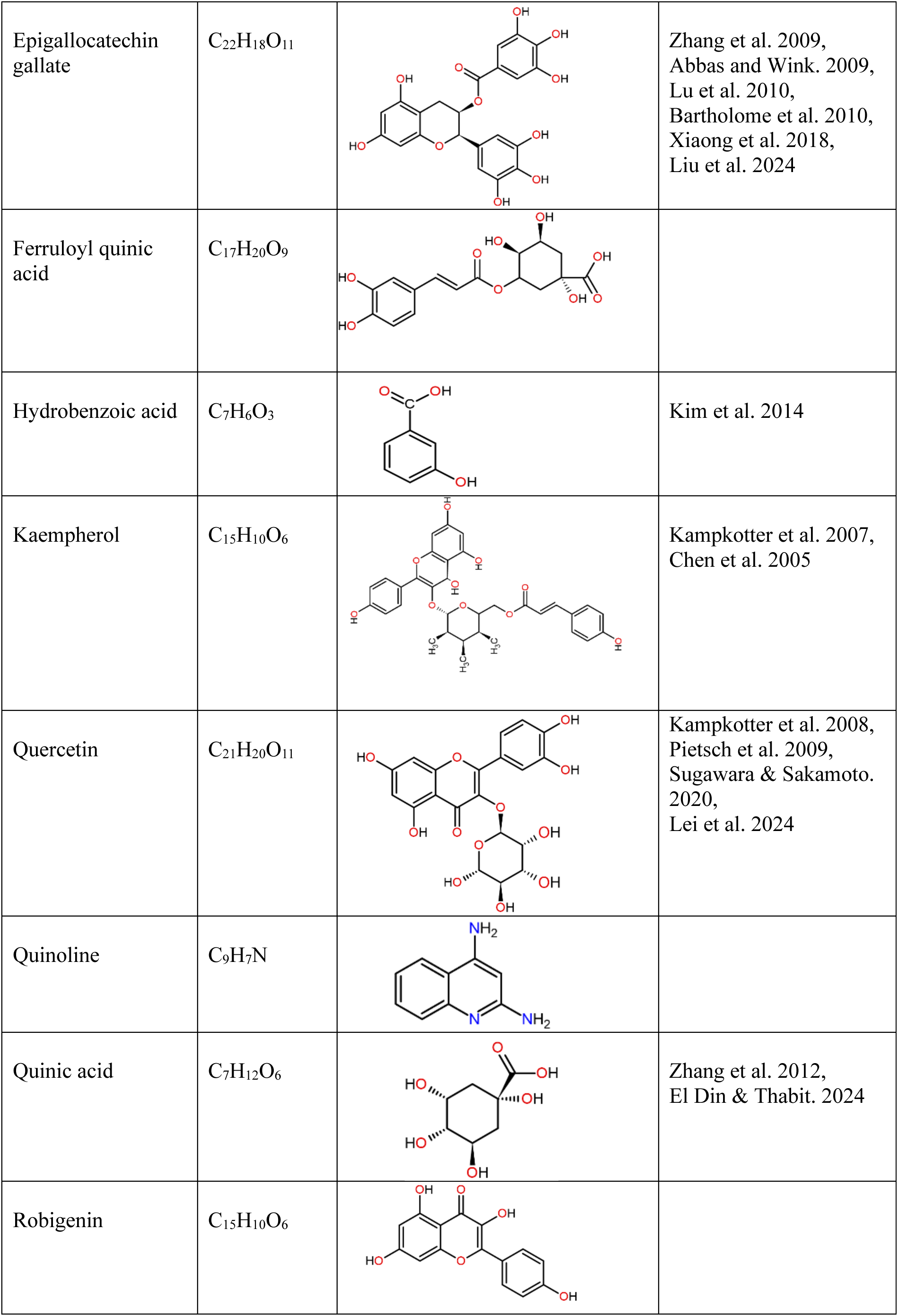
2D representation of phytochemicals structure.

| Compound | Chemical formula | 2D structure | References |
| --- | --- | --- | --- |
| Chlorogenic acid | C<sub>16</sub>H<sub>18</sub>O<sub>9</sub> |  | Zheng et al. 2017,<br>Sisawanto et al. 2022,<br>Cho et al. 2023 |
| Caffeic acid | C<sub>9</sub>H<sub>8</sub>O<sub>4</sub> |  | Li et al. 2021,<br>Gutierrez-Zetina et al.<br>2021, Xio et al. 2024 |
| Catechin | C<sub>15</sub>H<sub>14</sub>O<sub>6</sub> |  | Saul et al. 2009 |
| Coumaric acid | C<sub>9</sub>H<sub>8</sub>O<sub>3</sub> |  | Yue et al. 2019 |
| Coumaroyl quinic acid | C<sub>16</sub>H<sub>18</sub>O<sub>8</sub> |  |  |
| Curcumin | C<sub>21</sub>H<sub>20</sub>O<sub>6</sub> |  | Liao et al. 2011,<br>Yue et al. 2021,<br>Xu et al. 2023,<br>Savova et al. 2025 |
| Epicatechin | C<sub>15</sub>H<sub>14</sub>O<sub>6</sub> |  | González-Manzano et<br>al. 2012,<br>Duran et al. 2019 |
| Epigallocatechin gallate | $C_{22}H_{18}O_{11}$ | | Zhang et al. 2009,<br>Abbas and Wink. 2009,<br>Lu et al. 2010,<br>Bartholome et al. 2010,<br>Xiaong et al. 2018,<br>Liu et al. 2024 |
| Ferruloyl quinic acid | $C_{17}H_{20}O_9$ | | |
| Hydrobenzoic acid | $C_7H_6O_3$ | | Kim et al. 2014 |
| Kaempferol | $C_{15}H_{10}O_6$ | | Kampkotter et al. 2007,<br>Chen et al. 2005 |
| Quercetin | $C_{21}H_{20}O_{11}$ | | Kampkotter et al. 2008,<br>Pietsch et al. 2009,<br>Sugawara & Sakamoto.<br>2020,<br>Lei et al. 2024 |
| Quinoline | $C_9H_7N$ | | |
| Quinic acid | $C_7H_{12}O_6$ | | Zhang et al. 2012,<br>El Din & Thabit. 2024 |
| Robigenin | $C_{15}H_{10}O_6$ | | |

**Supplementary Figure 1.**
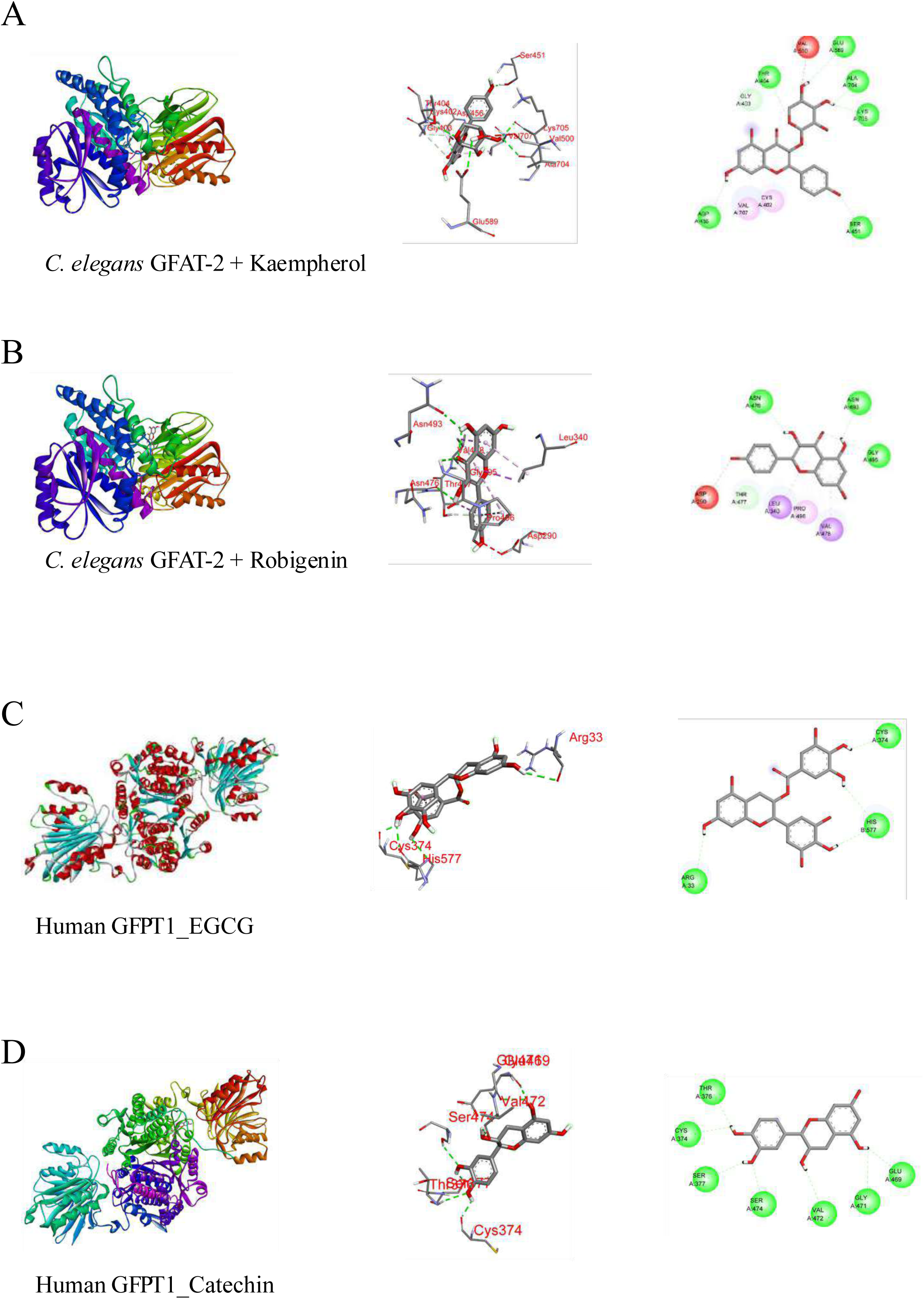
*In silico* interaction of phytochemicals with *C. elegans* GFAT-2 and human GFPT1. (A) Molecular docking of *C. elegans* GFAT-2 with Kaempherol. (B) Molecular docking of *C. elegans* GFAT-2 with Robigenin. (C) Molecular docking of human GFPT1 with Epigallocatechin gallate. (D) Molecular docking of human GFPT1 with Catechin.

## References

Labbadia J, Morimoto RI (2015). The biology of proteostasis in aging and disease. Annual Review of Biochem 84: 435–464. 10.1146/annurev-biochem-060614-033955

Klaips CL, Jayaraj GG, Hartl FU (2018) Pathways of cellular proteostasis in aging and disease. Journal of Cell Biol 217(1): 51–63. 10.1083/jcb.201709072

Taylor, R. C., & Dillin, A. (2011). Aging as an event of proteostasis collapse. Cold Spring Harbor Perspectives in Biology, 3(5), a004440. 10.1101/cshperspect.a004440

Hipp, M. S., Kasturi, P., & Hartl, F. U. (2019). The proteostasis network and its decline in ageing. Nature Reviews Molecular Cell Biology, 20(7), 421–435. 10.1038/s41580-019-0101-y

Chiti F, Dobson CM (2017) Protein Misfolding, Amyloid Formation, and Human Disease: A Summary of Progress Over the Last Decade. Annu Rev Biochem 86:27–68. doi:10.1146/annurev-biochem-061516-045115

Wilson DM 3rd, Cookson MR, Van Den Bosch L, Zetterberg H, Holtzman DM, Dewachter I (2023) Hallmarks of neurodegenerative diseases. Cell 186(4):693–714. doi:10.1016/j.cell.2022.12.032

Mack HID, Heimbucher T, Murphy CT (2018) The nematode Caenorhabditis elegans as a model for aging research. Drug Discovery Today: Disease Models 27: 3–13. 10.1016/j.ddmod.2018.11.001

Vimal P, Verma D, Bhunia PK, Kasturi P (2026) Caenorhabditis elegans as a Model Organism for Proteinopathies. In: Kumar, P., Kothari, V. (eds) Model Organisms in Biological Research. Springer, Singapore. 10.1007/978-981-92-0038-2_9

David DC, Ollikainen N, Trinidad JC, Cary MP, Burlingame AL, Kenyon C (2010) Widespread protein aggregation as an inherent part of aging in C. elegans. PLoS Biol 8(8):e1000450. doi: 10.1371/journal.pbio.1000450.

Reis-Rodrigues P, Czerwieniec G, Peters TW, Evani US, Alavez S, Gaman EA, Vantipalli M, Mooney SD, Gibson BW, Lithgow GJ, Hughes RE (2012) Proteomic analysis of age-dependent changes in protein solubility identifies genes that modulate lifespan. Aging Cell 11(1):120–7. doi: 10.1111/j.1474-9726.2011.00765.x.

Walther DM, Kasturi P, Zheng M, Pinkert S, Vecchi G, Ciryam P, Morimoto RI, Dobson CM, Vendruscolo M, Mann M, Hartl FU (2015) Widespread Proteome Remodeling and Aggregation in Aging C. elegans. Cell: 161(4):919–932. doi:10.1016/j.cell.2015.03.032

Narayan V, Ly T, Pourkarimi E, Murillo AB, Gartner A, Lamond AI, Kenyon C (2016) Deep Proteome Analysis Identifies Age-Related Processes in C. elegans. Cell Syst 3(2):144–159. doi: 10.1016/j.cels.2016.06.011

Zhu TY, Li ST, Liu DD, Zhang X, Zhou L, Zhou R, Yang B (2024) Single-worm quantitative proteomics reveals aging heterogeneity in isogenic Caenorhabditis elegans. Aging Cell 23(3):e14055. doi: 10.1111/acel.14055

Ciryam P, Tartaglia GG, Morimoto RI, Dobson CM, Vendruscolo M (2013) Widespread aggregation and neurodegenerative diseases are associated with supersaturated proteins. Cell Rep 5(3):781–90. doi: 10.1016/j.celrep.2013.09.043.

Chen JC, Wang R, Wei CC (2024) Anti-aging effects of dietary phytochemicals: From Caenorhabditis elegans, Drosophila melanogaster, rodents to clinical studies. Crit Rev Food Sci Nutr 64(17):5958–5983. doi: 10.1080/10408398.2022.2160961.

Cuanalo-Contreras K, Moreno-Gonzalez I (2019) Natural Products as Modulators of the Proteostasis Machinery: Implications in Neurodegenerative Diseases. Int J Mol Sci. 20(19):4666. doi: 10.3390/ijms20194666

Dhouafli Z, Cuanalo-Contreras K, Hayouni EA, Mays CE, Soto C, Moreno-Gonzalez I (2018) Inhibition of protein misfolding and aggregation by natural phenolic compounds. Cell Mol Life Sci 75(19):3521–3538. doi: 10.1007/s00018-018-2872-2.

Henríquez G, Gomez A, Guerrero E, Narayan M (2020) Potential Role of Natural Polyphenols against Protein Aggregation Toxicity: In Vitro, In Vivo, and Clinical Studies. ACS Chem Neurosci 11(19):2915–2934. doi: 10.1021/acschemneuro.0c00381.

Davinelli S, Medoro A, Hu FB, Scapagnini G (2025) Dietary polyphenols as geroprotective compounds: From Blue Zones to hallmarks of ageing. Ageing Res Rev. 108:102733. doi: 10.1016/j.arr.2025.102733.

Ren K, Lu D, Wang L, Yang J, Zhang H (2026) Phytochemicals modulating HSF-1-associated pathways: A systematic review of longevity-extending mechanisms in Caenorhabditis elegans. Ageing Res Rev. 121:103265. doi: 10.1016/j.arr.2026.103265.

Wedel S, Manola M, Cavinato M, Trougakos IP, Jansen-Dürr P (2018) Targeting Protein Quality Control Mechanisms by Natural Products to Promote Healthy Ageing. Molecules 23(5):1219. doi: 10.3390/molecules23051219.

Freyssin A, Page G, Fauconneau B, Rioux Bilan A (2018) Natural polyphenols effects on protein aggregates in Alzheimer’s and Parkinson’s prion-like diseases. Neural Regen Res.13(6):955–961. doi: 10.4103/1673-5374.233432.

Denzel MS, Storm NJ, Gutschmidt A, Baddi R, Hinze Y, Jarosch E, Sommer T, Hoppe T, Antebi A (2014) Hexosamine pathway metabolites enhance protein quality control and prolong life. Cell 156(6):1167–1178. doi: 10.1016/j.cell.2014.01.061.

Horn M, Denzel SI, Srinivasan B, Allmeroth K, Schiffer I, Karthikaisamy V, Miethe S, Breuer P, Antebi A, Denzel MS (2020) Hexosamine Pathway Activation Improves Protein Homeostasis through the Integrated Stress Response. iScience 23(3):100887. doi: 10.1016/j.isci.2020.100887.

Guergueltcheva V, Müller JS, Dusl M, Senderek J, Oldfors A, Lindbergh C, et al (2012) Congenital myasthenic syndrome with tubular aggregates caused by GFPT1 mutations. J Neurol 259(5):838–50. doi: 10.1007/s00415-011-6262-z.

Kroef V, Ruegenberg S, Horn M, Allmeroth K, Ebert L, Bozkus S, Miethe S, Elling U, Schermer B, Baumann U, Denzel MS (2022) GFPT2/GFAT2 and AMDHD2 act in tandem to control the hexosamine pathway. Elife 11:e69223. doi: 10.7554/eLife.69223.

Klappan AK, Hones S, Mylonas I, Brüning A (2012) Proteasome inhibition by quercetin triggers macroautophagy and blocks mTOR activity. Histochem Cell Biol 137(1):25–36. doi: 10.1007/s00418-011-0869-0.

Wang WW, Han R, He HJ, Li J, Chen SY, Gu Y, Xie C (2021) Administration of quercetin improves mitochondria quality control and protects the neurons in 6-OHDA-lesioned Parkinson’s disease models. Aging (Albany NY) 13(8):11738–11751. doi: 10.18632/aging.202868.

Zhou S, Qingman X, Zhang W, Zhang D, Liu S (2025) Quercetin as a Multi-Target Natural Therapeutic in Aging-Related Diseases: Systemic Molecular and Cellular Mechanisms. Phytother Res 39(10):4821–4869. doi: 10.1002/ptr.70078.

Ayuda-Durán B, González-Manzano S, Miranda-Vizuete A, Sánchez-Hernández E, R Romero M, Dueñas M, Santos-Buelga C, González-Paramás AM (2019) Exploring Target Genes Involved in the Effect of Quercetin on the Response to Oxidative Stress in Caenorhabditis elegans. Antioxidants (Basel) 8(12):585. doi: 10.3390/antiox8120585.

Sugawara T, Sakamoto K (2020) Quercetin enhances motility in aged and heat-stressed Caenorhabditis elegans nematodes by modulating both HSF-1 activity, and insulin-like and p38-MAPK signalling. PLoS One 15(9):e0238528. doi: 10.1371/journal.pone.0238528.

Yu KH, Lee CI (2020) Quercetin Disaggregates Prion Fibrils and Decreases Fibril-Induced Cytotoxicity and Oxidative Stress. Pharmaceutics 12(11):1081. doi: 10.3390/pharmaceutics12111081.

Ge, S. X., Jung, D., & Yao, R. (2020). ShinyGO: A graphical gene-set enrichment tool for animals and plants. Bioinformatics, 36(8), 2628–2629. 10.1093/bioinformatics/btz931

Thompson, J. D., Higgins, D. G., & Gibson, T. J. (1994). CLUSTAL W: Improving the sensitivity of progressive multiple sequence alignment through sequence weighting, position-specific gap penalties and weight matrix choice. Nucleic Acids Research, 22(22), 4673–4680. 10.1093/nar/22.22.4673

Gouet, P., Courcelle, E., Stuart, D. I., & Métoz, F. (1999). ESPript: Analysis of multiple sequence alignments in PostScript. Bioinformatics, 15(4), 305–308. 10.1093/bioinformatics/15.4.305

Abramson, J., Adler, J., Dunger, J., Evans, R., Green, T., Pritzel, A., Ronneberger, O., Willmore, L., Ballard, A. J., Bambrick, J., Bodenstein, S. W., Evans, D. A., Hung, C.-C., O’Neill, M., Reiman, D., Tunyasuvunakool, K., Wu, Z., Žemgulytė, A., Arvaniti, E., … Jumper, J. (2024). Accurate structure prediction of biomolecular interactions with AlphaFold 3. Nature, 630(8016), 493–500. 10.1038/s41586-024-07487-w

Berman, H. M., Westbrook, J., Feng, Z., Gilliland, G., Bhat, T. N., Weissig, H., Shindyalov, I. N., & Bourne, P. E. (2000). The Protein Data Bank. Nucleic Acids Research, 28(1), 235–242. 10.1093/nar/28.1.235

Dassault Systèmes. (2025). BIOVIA Discovery Studio Visualizer [Computer software]. Dassault Systèmes.

Pettersen, E. F., Goddard, T. D., Huang, C. C., Meng, E. C., Couch, G. S., Croll, T. I., Morris, J. H., & Ferrin, T. E. (2021). UCSF ChimeraX: Structure visualization for researchers, educators, and developers. Protein Science: A Publication of the Protein Society, 30(1), 70–82. 10.1002/pro.3943

Kim, S., Chen, J., Cheng, T., Gindulyte, A., He, J., He, S., Li, Q., Shoemaker, B. A., Thiessen, P. A., Yu, B., Zaslavsky, L., Zhang, J., & Bolton, E. E. (2019). PubChem 2019 update: Improved access to chemical data. Nucleic Acids Research, 47(D1), D1102–D1109. 10.1093/nar/gky1033

Daina, A., Michielin, O., & Zoete, V. (2017). SwissADME: A free web tool to evaluate pharmacokinetics, drug-likeness and medicinal chemistry friendliness of small molecules. Scientific Reports, 7(1), Article 42717. 10.1038/srep42717

Gu, Y., Yu, Z., Wang, Y., Chen, L., Lou, C., Yang, C., Li, W., Liu, G., & Tang, Y. (2024). admetSAR3.0: A comprehensive platform for exploration, prediction and optimization of chemical ADMET properties. Nucleic Acids Research, 52(W1), W432–W438. 10.1093/nar/gkae298

PerkinElmer Informatics. (2003). ChemDraw Pro 8.0 [Computer software]. PerkinElmer Informatics.

Dallakyan, S., & Olson, A. J. (2015). Small-molecule library screening by docking with PyRx. Methods in Molecular Biology, 1263, 243–250. 10.1007/978-1-4939-2269-7_19

